# Global Structure of the Intrinsically Disordered Protein Tau Emerges from its Local Structure

**DOI:** 10.1101/2021.11.23.469691

**Authors:** Lukas S. Stelzl, Lisa M. Pietrek, Andrea Holla, Javier Oroz, Mateusz Sikora, Jürgen Köfinger, Benjamin Schuler, Markus Zweckstetter, Gerhard Hummer

**Affiliations:** Department of Theoretical Biophysics, Max Planck Institute of Biophysics, Max-von-Laue-Straße 3, 60438 Frankfurt am Main, Germany; Faculty of Biology, Johannes Gutenberg University Mainz, Gresemundweg 2, 55128 Mainz, Germany; KOMET 1, Institute of Physics, Johannes Gutenberg University Mainz, 55099 Mainz, Germany; and Institute of Molecular Biology (IMB), 55128 Mainz, Germany; Department of Biochemistry, University of Zurich, 8057 Zurich, Switzerland; German Center for Neurodegenerative Diseases (DZNE), von-Siebold-Str. 3a, 37075 Göttingen, Germany. 2; Faculty of Physics, University of Vienna, Kolingasse 14-16, 1090 Vienna, Austria; Department of Physics, University of Zurich, 8057 Zurich, Switzerland; Department for NMR-based Structural Biology, Max Planck Institute for Biophysical Chemistry, Am Faßberg 11, 37077 Göttingen, Germany; Institute for Biophysics, Goethe University Frankfurt, Max-von-Laue-Straße 9, 60438 Frankfurt am Main, Germany

## Abstract

The paradigmatic disordered protein tau plays an important role in neuronal function and neurodegenerative diseases. To disentangle the factors controlling the balance between functional and disease-associated conformational states, we build a structural ensemble of the tau K18 fragment containing the four pseudorepeat domains involved in both microtubule binding and amyloid fibril formation. We assemble 129-residue-long tau K18 chains at atomic resolution from an extensive fragment library constructed with molecular dynamics simulations. We introduce a reweighted hierarchical chain growth (RHCG) algorithm that integrates experimental data reporting on the local structure into the assembly process in a systematic manner. By combining Bayesian ensemble refinement with importance sampling, we obtain well-defined ensembles and overcome the problem of exponentially varying weights in the integrative modeling of long-chain polymeric molecules. The resulting tau K18 ensembles capture nuclear magnetic resonance (NMR) chemical shift and J-coupling measurements. Without further fitting, we achieve excellent agreement with measurements of NMR residual dipolar couplings. The good agreement with experimental measures of global structures such as singlemolecule Förster resonance energy transfer (FRET) efficiencies is improved further by ensemble refinement. By comparing wild-type and mutant ensembles, we show that pathogenic single-point P301 mutations shift the population from the turn-like conformations of the functional microtubule-bound state to the extended conformations of disease-associated tau fibrils. RHCG thus provides us with an atomically resolved view of the population equilibrium between functional and aggregation-prone states of tau K18, and demonstrates that global structural characteristics of this intrinsically disordered protein emerge from its local structure.

## Introduction

Intrinsically disordered proteins (IDPs) are enriched in the proteomes of higher eukaryotes, where they perform essential functions.^1–3^ In healthy neurons, the paradigmatic IDP tau binds and stabilizes microtubules.^1^ In diseased neurons, tau loses the ability to bind to microtubules and forms the toxic aggregates associated with Alzheimer’s and other neurodegenerative diseases.^2^ Hyperphosphorylation of tau correlates with the progression of Alzheimer’s disease. Tau has recently been shown to form biomolecular condensates.^3–6^ Dysregulation of the formation of biomolecular condensates by mutations^7^ and aberrant post-translational modifications such as phosphorylation^4,7^ may underlie the pathogenicity of tau. Some tau mutations, e.g., P301L and P301S, show drastic effects in patients and are used in mouse models of tau pathology.^8,9^ The conformational dynamics of tau around P301 may play a direct role in modulating the aggregation of tau in disease,^10–12^ as studied also by molecular dynamics (MD) simulations of tau fragments.^12^ Efforts to gain a clearer picture of the local conformational dynamics of tau promise a deeper understanding of its roles in health and disease.

The challenges in resolving structural ensembles of IDPs ask for an integrative approach.^13^ Important progress in dealing with the high flexibility of disordered biomolecules has been made using nuclear magnetic resonance (NMR) spectroscopy,^14–17^ solution X-ray scattering (SAXS)^18^ and single-molecule Förster resonance energy transfer (FRET).^19–23^ To harness the full power of these experiments and interpret the data in detail, the construction of ensembles of structures ^24–32^ has proved to be a powerful strategy, especially for the interpretation of NMR experiments and the combination of multiple experimental methods. ^31,33,34^ For instance, Borgia et al.^32^ combined data from single-molecule FRET, SAXS, dynamic light scattering, and fluorescence correlation spectroscopy with MD simulations to characterize the ensembles of a marginally stable spectrin domain and the IDP ACTR over a broad range of solution conditions. Gomes and co-workers^35^ recently described an ensemble of the disordered N-terminal region of the Sic protein, obtained by integrating different combinations of SAXS, single-molecule FRET and NMR experiments using the ENSEMBLE approach. ^36^

Structural ensembles obtained from computational modeling can be combined with experimental data by using Bayesian and maximum entropy ensemble refinement methods. ^29,37-45^ The Bayesian formulation accounts naturally for uncertainties in the measurements, the model used to generate the ensemble, and the calculation of observables from the ensemble members.^39^ Input ensembles^46^ are obtained, e.g., from MD simulations^44,47-49^ or chain growth, ^26,28,50-53^ and are then minimally modified to account for the experimental observations. However, for long protein or nucleic acid chains, it is difficult to create initial ensembles that have sufficient overlap with the true ensemble for reliable ensemble refinement. For experimental data that report on the local structure along the chain of a disordered protein, we expect that cumulative systematic errors in the MD force field will cause the summed squared error *χ*^2^ between model and experiment to grow linearly with the length of the chain. As a consequence, the overlap between input and true ensemble deteriorates exponentially as the chain grows in length. Consequently, for long IDPs only a few chains will tend to dominate the ensemble after refinement, with the rest of the large ensemble being mostly irrelevant.

The problem of poor overlap between initial and true ensemble can be overcome by applying a bias already in the generation of the initial ensemble, e.g., by imposing restraints directly on observables or related quantities in the initial MD simulations. In an early combination of biased chain growth with Bayesian weighting applied to tau K18,^28^ overlapping peptide fragments were stitched together. Fragment selection was biased to double the radius of gyration in an otherwise overly compact ensemble. Steric clashes were resolved by energy minimization in implicit solvent, and high-energy structures were randomly removed in a pruning step. Excellent agreement with NMR observables^27^ could be achieved by adjusting the weights of the ensemble members. However, formal and practical questions are raised: how does one incorporate experimental data already during chain growth without compromising the Bayesian framework of ensemble refinement, where such information would normally be used a posteriori? And how does one ensure that the final ensemble is well defined and fully reproducible?

We will show here that in a Bayesian formulation any bias in ensemble generation can be accounted for fully and quantitatively in a final global refinement step by exploiting the direct connection of ensemble refinement to traditional free energy calculations. ^39^ Meaningful input ensembles can thus be generated without sacrificing the rigor and reproducibility of the ensemble refinement procedure.

We propose reweighted hierarchical chain growth (RHCG) as a general method to integrate data reporting on local structure into models of disordered and flexible polymeric molecules such as disordered proteins or nucleic acids. Protein chains are assembled from fragment structures, as obtained here from MD simulations. As in hierarchical chain growth (HCG), ^52^ chains with steric clashes are consistently removed in such a way that the resulting ensemble does not depend on arbitrary choices such as the direction of chain growth, N-to-C versus C-to-N. In RHCG, fragment choice is biased according to experiments reporting on the local structure. In a final reweighting step, any resulting bias is then removed. RHCG is thus a form of importance sampling.

Using RHCG, we arrive at an integrative model of tau K18 with atomic resolution. Tau K18 contains the four pseudorepeat domains R1-R4 involved both in functional binding to microtubules^54^ and in forming amyloid fibrils.^10,12^ NMR chemical shift data that report on local structure are incorporated already during chain growth. We then show that these ensembles also capture the global structure of tau K18, as probed by NMR, RDC, singlemolecule FRET and SAXS measurement.

By comparing wild-type (WT) and mutant sequences, we provide a molecular view of possible differences between tau in a healthy cell and tau with pathogenic mutations. Our modeling of tau K18 reveals turns as in microtubule-bound states and extended structures as in tau fibrils. We found that pathogenic single-point P301 mutations shift the equilibrium from the former to the latter, emphasizing the close connection between functional forms of tau in solution and the fibrillar structures in tau-associated pathologies.

## Theory

### Bayesian Ensemble Refinement of Polymeric Molecules

We combine molecular simulations with ensemble refinement to create ensembles of proteins or nucleic acids that faithfully reflect the distribution of conformations in experiment. To create an initial ensemble, we adapt the hierarchical chain growth (HCG) method introduced recently,^52^ as described in detail below. We then use Bayesian Inference of Ensembles (BioEn)^39^ to adjust the weights of the individual ensemble members according to the experimental data, e.g., NMR chemical shifts.

BioEn ensemble refinement minimally adjusts the vector ***w*** = (*w*_1_,…,*w_C_*) of normalized weights of individual chains *c* = 1,…,*C* in the ensemble to match the experimental data. We define a posterior *P*(***w***|data, *I*) as a function of the weights ***w***,

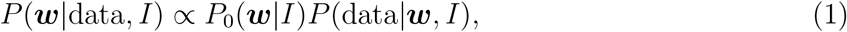

with *P*_0_(***w**|I*) the prior and *P*(data|***w**,I*) the likelihood. Here, *I* denotes background information, e.g., that we model polymeric molecules with internal structure. The BioEn maximum-entropy prior^38^ is given by

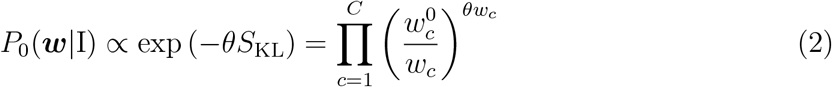

*θ* is a hyperparameter that controls the strength of the entropy regularization and thus expresses our confidence in the initial ensemble of chains.^39^ *S*_KL_ is the Kullback-Leibler (KL) divergence

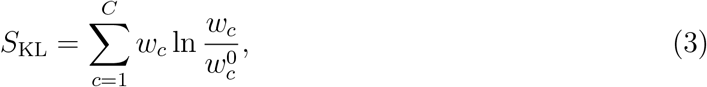

which reports how close the normalized refined weights *w_c_* are to the normalized reference weights 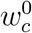.

Assuming Gaussian uncorrelated errors, the likelihood is *P*(data|***w**, I*) ∝ exp(−*χ*^2^/2) with

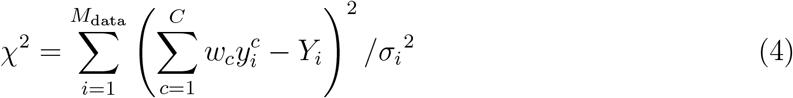

The first sum is over the different experimental observations *i* = 1,…,*M*_data_ with measured values *Y_i_*, and the second sum over the ensemble members *c* = 1,…,*C*. For each chain *c* and observable *i*, we use a forward model to compute individual observations 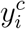. The error 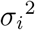 is the sum of the squared standard errors of the measurements *Y_i_* and the forward calculations 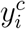.

In applications of BioEn to long biopolymers, small but systematic weight corrections at the monomer level can add up to large corrections overall. For NMR chemical shifts, for instance, the sum over *i* in eq 4 corresponds to a sum over residues. As a result, the *χ*^2^ statistic is extensive, i.e., it tends to grow linearly with the length of the chain. Reweighting of assembled chains thus becomes progressively more challenging as the length of the chain grows (i.e., for chains with more fragments). The reason is that it becomes progressively unlikely that all fragments in an assembled chain occupy the relevant subspace with proper weight. As a result, chains will contribute with very uneven weights after BioEn reweighting. In other words, a few chains will dominate, and the rest of the large ensemble is more or less irrelevant.

### Reweighted Hierarchical Chain Growth

We address the problem of poor overlap between initial and true ensemble by using importance sampling. In MD simulations of complete biopolymer chains, bias potentials could be introduced, acting for instance on the torsion angles to better match NMR chemical shifts or J-couplings. Here, we focus instead on fragment-based chain growth. The key idea is to grow chains by using fragment libraries that have already been biased to enrich the ensemble with members of high weight, and then to correct for this biased choice of fragments in a final reweighting step. If the bias weights were chosen perfectly, the final step would give each chain equal weight.

In RHCG, we adapt HCG^52^ to assemble polymer chains from fragments. At each of the *N* positions, fragments are picked at random from a fragment library and then combined by superimposition of residues at their termini with the equivalent residues in the adjacent fragments. Any models with steric clashes are discarded. In HCG, all fragments have equal weight; in RHCG, the fragments in the library 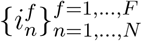 (with *F* the number of fragments created at position *n*) are picked according to a weight 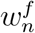 normalized to 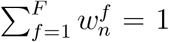 for all *n*. These weights are our initial guess as to how likely a particular fragment is in the final reweighted ensemble of chains. The probability *p*[***f**^c^*] for a particular chain *c* to be created in this way is given by the product of weights for each of its fragments,

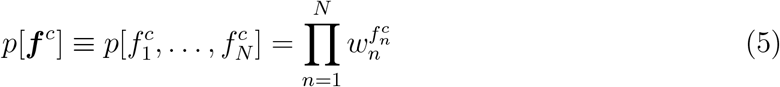

where 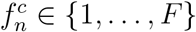 is the index of fragment *n* in chain *c*.

### BioEn Reweighting of Assembled Chains

After the biased assembly of an ensemble chains, we use BioEn^39,40^ to correct for the bias in chain growth and to reweight the entire ensemble globally. To correct for the bias in chain assembly, chain *c* enters the global BioEn refinement with a relative weight proportional to the reciprocal of the bias probability, 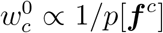, with which its fragments were selected. Normalization of these relative weights gives us

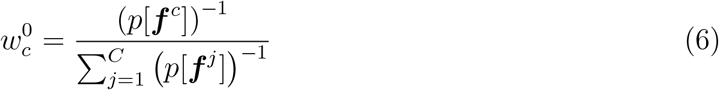

or, expressed more compactly in terms of reciprocal weight factors,

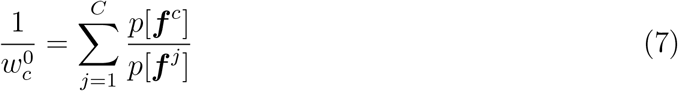

where the sum extends over the *C* chains of the ensemble. To the ensemble with these initial weights we then apply BioEn reweighting, using as reference experimental data reporting on local or global structural properties.

### Chain Growth with Non-Bonded Interactions beyond Steric Repulsion

Fragment assembly can, in principle, be extended to account for non-bonded interactions beyond steric repulsion to account, e.g., for electrostatic interactions between fragments. ^55^ This can be accomplished by using a free energy function 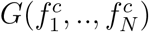 that describes the inter-fragment interactions in chain *c* and can be calculated from an implicit solvent model or, by free energy calculations, from explicit solvent models. Chains *c* assembled from fragments 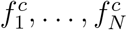 are then weighted by an additional factor 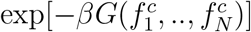 with 1/*β* = *k_B_T* and *k_B_* the Boltzmann constant and *T* the absolute temperature. In the Bayesian formulation, the normalized reference weight of chain *c* in an ensemble of *C* chains then becomes

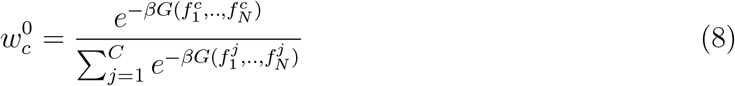

To sample efficiently from this distribution, one can again use importance sampling by performing hierarchical assembly^52^ with biased fragment selection. If, as above, 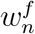 is the bias weight factor to choose fragment *f* at position *n*, then eq 7 becomes

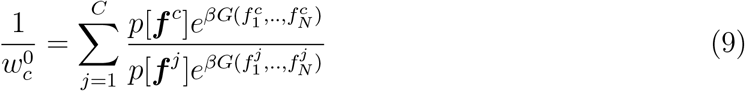

Here, we use only excluded volume interactions, which amounts to exp(−*G*) = 1 for chains without inter-fragment steric clashes and exp(−*G*) = 0 with clashes.

### Assessment of Importance Sampling

In ideal importance sampling, we would grow chains of equal relative importance. Global BioEn reweighting would then give each member of the resulting ensemble equal weight, *w_c_* = 1/*C*. We use the KL divergence of the BioEn-optimized weights *w_c_* from ideal importance sampling to assess the effectiveness of our bias in chain growth:

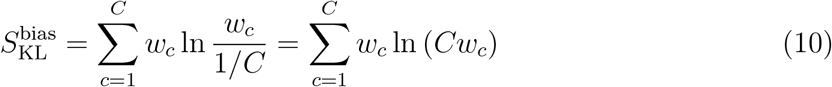

If 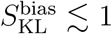, the overlap between the ensembles produced by biased chain growth and after BioEn refinement is large; conversely, if 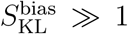, the chain growth protocol should be optimized.

## Methods

### Hierarchical and Reweighted Hierarchical Chain Growth

We generated structural ensembles of tau K18 (residues 244-372) using HCG^52^ and RHCG. Tau structures were assembled from 43 pentamer fragments with two residues overlap between subsequent fragments. All fragments had their N and C termini capped by acetyl and N-methyl groups, respectively. The first (N-terminal) fragment started from the last residue outside tau K18, which was then removed in chain assembly. Fragment structures were sampled in all-atom replica exchange molecular dynamics (REMD) with explicit solvent. For each fragment, we used 24 replicas spanning a temperature range of 278-420 K. Each pentamer fragment was simulated for 100 ns as in our previous study.^52^ We used structures from the *T* = 278 K ensemble to assemble tau K18 chains, which corresponds to the temperature of the NMR experiments.^27^ To investigate the effect of point mutations at the P301 position, we also sampled fragments with P301 and mutations P301L, P301S and P301T. We repeated fragment simulations for WT P301, P301L, P301S and P301T fragments with residue 301 at the central position of their respective fragments instead of the second position of its respective pentameter. We note that in all fragment REMD simulations P301 was sampled exclusively as trans isomer.

We biased the fragment selection in RHCG according to C_*α*_ chemical shifts measured by NMR. At each fragment position *n*, we performed independent BioEn reweighting^39,40^ using the chemical shift data reported for the non-terminal residues in this fragment (Supporting Information (SI) Text). A large confidence parameter of *θ* =10 ensured improved consistency of the chemical shifts (with the average *χ*^2^ across fragments dropping from 0.856 to 0.688) with minimal weight changes (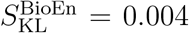 on average). These local BioEn calculations gave us fragment weight factors 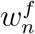.

We then used RHCG to build ensembles of between 2000 and 10^6^ WT tau K18 models from the reweighted fragment libraries. For reference, we also constructed unbiased ensembles of WT tau using HCG ^52^ with unweighted fragment libraries. HCG was also used to construct tau K18 ensembles of P301 mutants. If not specified otherwise, the results shown are for ensembles of *C* = 50000 chains. Following the procedure described in ref 52, we assembled 10000 representatives at each hierarchy level below the final assembly level to sample a high diversity of possible local conformations. At the final level, full-length models were assembled from this pool. The assembly process was trivially parallelized by using different random number seeds. In a final step, the RHCG ensembles were reweighted using BioEn to correct for the biased fragment choice while retaining consistency with the NMR chemical shift data. In this global BioEn reweighting step, the confidence parameter was set to *θ* = 5 according to an L-curve analysis (SI Text and Figure S1A). The resulting ensembles were structurally diverse and, among 50000 HCG and RHCG structures, did not contain any knots (SI Text).

### Calculation of Experimental Observables

#### NMR Secondary Chemical Shifts and J couplings

For comparison with NMR experiments, we calculated chemical shifts from fragments and full-length structures using SPARTA+.^56^ We subtracted random-coil shifts calculated using POTENCI^57^ to compare to secondary chemical shifts Δ*C*. We computed ^3^*J*_HNH*α*_ couplings with the Karplus parameters by Vögeli et al.^58^ with the mdtraj Python library.^59^

#### NMR Residual Dipolar Couplings

RDCs were calculated from the ensembles of fulllength structures with PALES^60,61^ in the steric alignment mode. The value 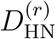 for a particular residue *r* was calculated by computing the alignment of each chain *c* in the ensemble with PALES and then taking the average over all structures

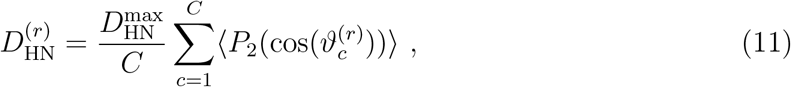

where 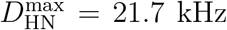 for an idealized amide bond length of 1.04 Å,^62^ 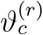 is the angle between the amide bond vector of residue *r* in chain *c* and the external magnetic field, *P*_2_(*x*) = (3*x*^2^ – 1)/2 is the second-order Legendre polynomial, and 〈···〉 denotes an average over the orientations of the chain biased by the alignment.

#### Small-Angle X-Ray Scattering

We used FoXS^63^ to calculate SAXS intensity profiles for the tau K18 structures in an ensemble, and then calculated the weighted average over the ensemble. In the FoXS calculations, we took the solvation shell into account by setting *c*_2_ = 3. The excluded-volume parameter was set to the default value of *c*_1_ = 1. Geometric *R_G_* values were computed using the MDAnalysis library.^64,65^ To compare measured scattering intensities to those predicted for the weighted ensemble, *I*(*q*)_sim_, we first estimated an intensity scale factor *a* and a constant for background correction *b* according to

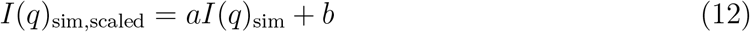

by performing least-square fitting. The best fit to experiment was achieved with the coefficients a = 1.11 × 10^−11^ and b = 3.75 × 10^−05^. We further took possible mild aggregation into account by approximating the scattering intensity of possible aggregates as

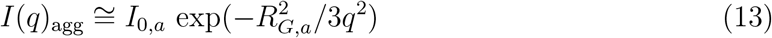

with *q* being the scattering vector. By adding *I*(*q*)_agg_ to *I*(*q*)_sim,scaled_ and adjusting the respective amplitudes as well as *R_G,a_* by least-square fitting, we accounted for the observed increase in scattering intensity at low *q*-values. A second set of scattering data^18^ is cut off at low *q* and, therefore, no correction is required.

#### Comparison to Single-Molecule FRET Experiments

We compared C_*α*_-C_*α*_ distances extracted from FRET experiments using the SAW-*ν* polymer model^66^ to RHCG models. To quantify the effect of the fluorescent dyes on the distance distribution, we performed additional calculations in which we adapted the RHCG method to add dyes^67^ during chain growth (SI Text and Figure S2).

### Experiments

#### Single-Molecule FRET Experiments

For the single-molecule FRET experiments, tau K18 was labeled with Alexa Fluor 488 and CF660R at C291 and C322 (SI Text). The labeled tau K18 was diluted to a concentration of 100 pM in 50 mM sodium phosphate buffer, pH 6.8, 1 mM DTT, 0.001% Tween 20 or 20 mM HEPES, 5 mM KCl, 10 mM MgCl_2_, pH 7.4, 1 mM DTT, 0.001% Tween 20. The experiments were performed at 295 K on a MicroTime 200 confocal single-molecule instrument (Pico-Quant, Berlin, Germany) as described in detail in the SI Text. The SAW-*ν* model was used to analyze the single-molecule FRET data to extract distances and the polymer properties of tau K18^66^ (SI Text).

#### Small-Angle X-Ray Scattering Experiments

SAXS data were collected at 298 K from monodisperse samples of K18 ranging from 50 *μ*M to 67 *μ*M in 20 mM Hepes, 5 mM KCl, 10 mM MgCl_2_, 1 mM DTT at pH 7.4. Scattering profiles were analyzed with standard procedures using ATSAS.^68^ SAXS measurements were performed at DESY (Hamburg, Germany) and Diamond Light Source (Oxford, UK) stations.

## Results and Discussion

### RHCG Produces a Diverse Ensemble of tau K18 Chains

During chain assembly, we applied a gentle bias on the fragment choice by using fragment weights from BioEn reweighting against C_*α*_ chemical shifts. To correct for the bias, the assembled chains were then reweighted with BioEn, again using the chemical shift data as experimental reference. In this global BioEn reweighting step, the chains were given near-uniform weights *w_c_* with 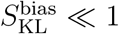 (Figure S1B). The resulting structural ensemble of tau is comprised of highly diverse structures at atomic resolution (Figure 1C). The typical C_*α*_ root-mean-square distance (RMSD) between two chains is about 26 Å (Figure S3 and SI Text).

**Figure 1:**
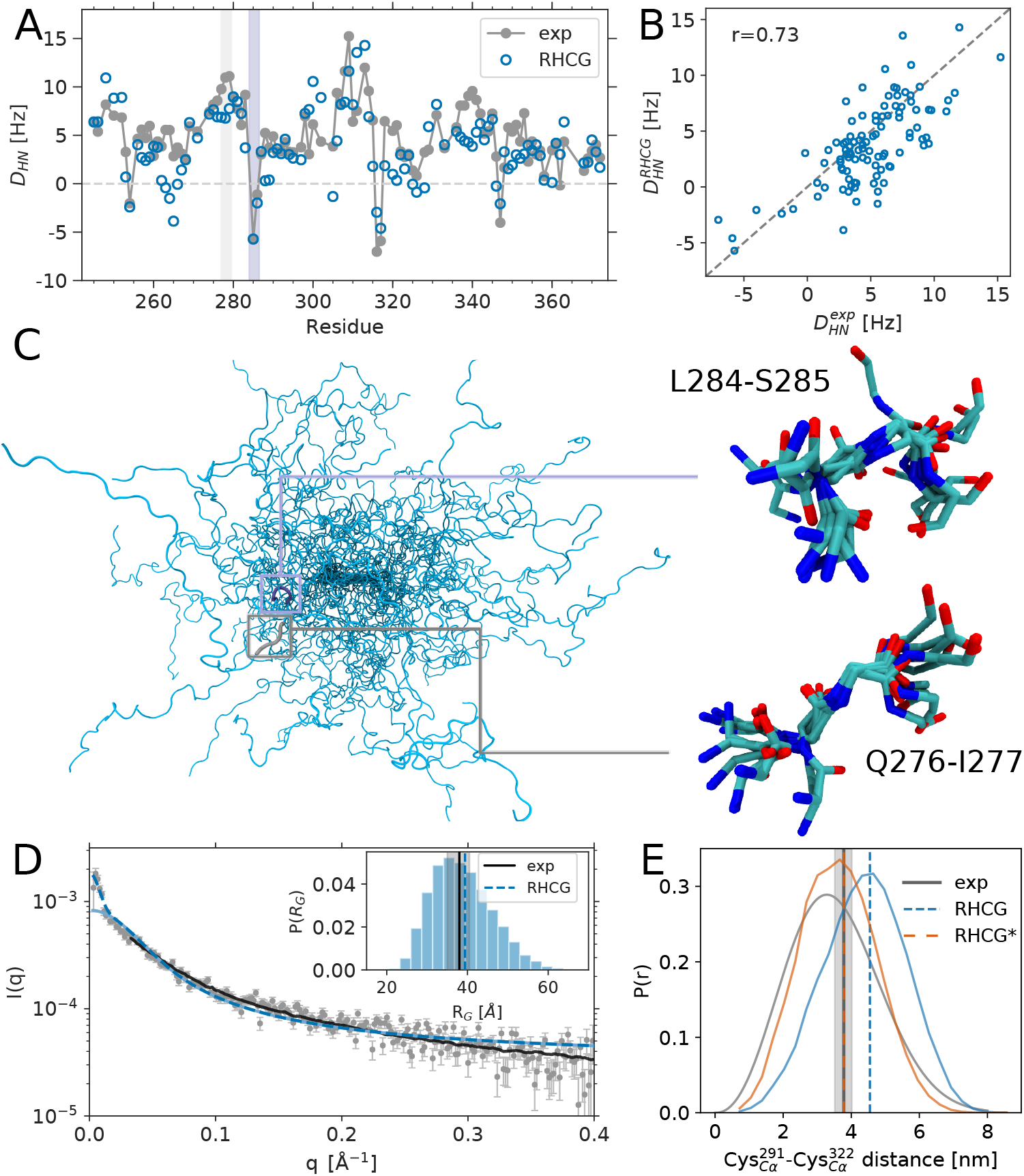
Atomic-resolution ensemble from RHCG reproduces global structural features of tau K18. (A) Comparison of experimental (grey) and predicted (blue) ^1^H-^15^N RDCs, which were not used in the construction of the RHCG ensemble (see Table S1 for the amino acid sequence of tau K18). (B) Scatter plot of calculated and measured RDCs. (C) Backbone traces of 30 members of the RHCG ensemble. Zoom-ins show superpositions of ten representative structures of a turn at position L284-S285 (top) and an extended segment at position Q276-I277 (bottom) with negative and positive RDCs, respectively, as highlighted by shading in panel A. (D) Comparison of calculated (blue) and experimental SAXS scattering intensity profiles (grey symbols) and from ref 18 (black line). Dashed line: Predicted scattering intensity *I*(*q*)_RHCG_ + *I*(*q*)_agg_ in the presence of mild aggregation. Inset: Distribution of *R_G_* in the RHCG ensemble. Vertical dashed lines indicate the average *R_G_* from RHCG (blue) and experiment^18^ (grey; ±SEM shown by shading). (E) Distribution of C*α*-C*α* FRET-label distance inferred from FRET experiments using the SAW-*ν* model^66^ (grey), RHCG (blue), and RHCG* (orange). Root-mean-square distances are indicated as (dashed) vertical lines.

### RHCG Models of tau K18 Capture the Average Local Structure of tau as Reported by NMR

Chemical shifts are accurate reporters of local structure and secondary structure.^16,17,27,29,56,69^ Overall, we found that the C_*α*_ chemical shifts calculated for the RHCG ensemble of tau K18 are close to random coil values, with secondary chemical shifts Δ*C* mostly close to zero. Despite the residual amplitude typically being smaller than the error of ≈1 ppm^56^ in the forward chemical shift calculation, the models capture important features of the variation of experimental secondary chemical shifts along the tau K18 amino acid sequence, such as a drop in secondary chemical shift going from L285 to V300. HCG without reweighting of the fragment library underestimates the populations of extended and *β*-strand like structures and overestimates the helical-like conformations. Going from HCG to RHCG, the average residual drops from 0.35 ppm to 0.27 ppm, and Pearson’s *r* for the secondary chemical shifts Δ*C* of the C_*α*_ atoms increases from 0.28 to 0.41. RHCG lowers in particular positive Δ*C* values, e.g., at the S420 position (Figure S4A,B). In light of the considerable uncertainties in the forward calculation (≈1 ppm) and the small Δ*C* amplitudes, a lower *θ* value resulting in an even tighter fit was not justified (Figure S1A).

We also calculated NMR ^3^*J*_HNH*α*_ couplings, which report primarily on the *ϕ*-dihedral angles of the protein backbone. The couplings calculated for our models agree well with the NMR experimental data^27^ (Figure S5). Also in terms of ^3^*J*_HNH*α*_, RHCG somewhat improves the representation of the local structures over HCG, as reflected by the increase of Pearson’s *r* from 0.59 to 0.62. Overall we conclude that reweighting in fragment assembly alleviates the small but systematic deviations caused by small imbalances in state-of-the-art force fields used to generate fragment libraries. As a result, the local structure of the tau K18 chains produced by RHCG is more consistent with NMR chemical shift and J-coupling experiments.

### The RHCG ensemble of tau K18 Reproduces the Experimental NMR Residual Dipolar Couplings

We calculated the RDCs for the assembled tau K18 chain using the steric alignment mode of PALES, ^61^ and then averaged the RDC values over the ensemble with the respective weight of the chain. The measured^27^ and calculated RDCs agree remarkably well and capture both the signature as a function of position along the chain (Figure 1A) and the magnitude at individual residue positions (Figure 1B). Without further fitting, we obtained Pearson *r* correlation coefficients of 0.73 for RHCG and 0.70 for HCG for tau K18 ensembles of 50000 models. This consistency not only validates the ensemble, but also gives direct insights into the interpretation of the RDCs measured for IDPs. RDCs inform on how restricted a chain is locally, with larger absolute RDCs expected for more restricted segments than for fully flexible segments.^15^ The RDC *D*_HN_ ∝ 〈*P*_2_(cos(*θ*))〉 reports on the relative orientation of an amide bond vector with respect to the magnetic field. Changes in the sign of the measured RDCs have been interpreted as changes in the direction of the protein backbone. ^27^ Our conformational ensemble reproduces the four changes in the sign of *D*_HN_ found in experiments.^27^ Importantly, as highlighted for the region centered on L284-S385 in Figure 1C, our structures on average trace a turn in the region where the sign changes, as indicated by a shortened distance across the four-residue segments (Figure S6). By contrast, in regions such as Q276-I277, where the sign of *D*_HN_ does not change, our structures do not show a preference in the chain direction and scatter around an average straight chain (Figure 1C). We note that simple polymeric models that ignore amino acid chemistry and the correlations between subsequent residues tend not to capture the trends in the experimental RDCs, as previously noted. ^15,27,70^

### Residual Dipolar Coupling Calculations Require Large Ensemble Sizes

The need for large ensembles has been highlighted before. ^26^ Building large ensembles relies on the possibility to quickly generate statistically independent atomic-resolution models of IDPs. The RDC values predicted for particular residues in our models are widely and asymmetrically distributed with a range of about ±25 Hz (Figure 2A). By contrast, the experimental average is roughly in the range of –5 to 10 Hz (Figure 1A). As a result, RDCs calculated from small ensembles are biased (Figure 2B). We found that relatively large ensembles of ≥ 10000 tau K18 chains are needed to get converged RDC values (Figure 2B). We found in particular that Pearson’s *r* correlation coefficient improved with increasing ensemble size.

**Figure 2:**
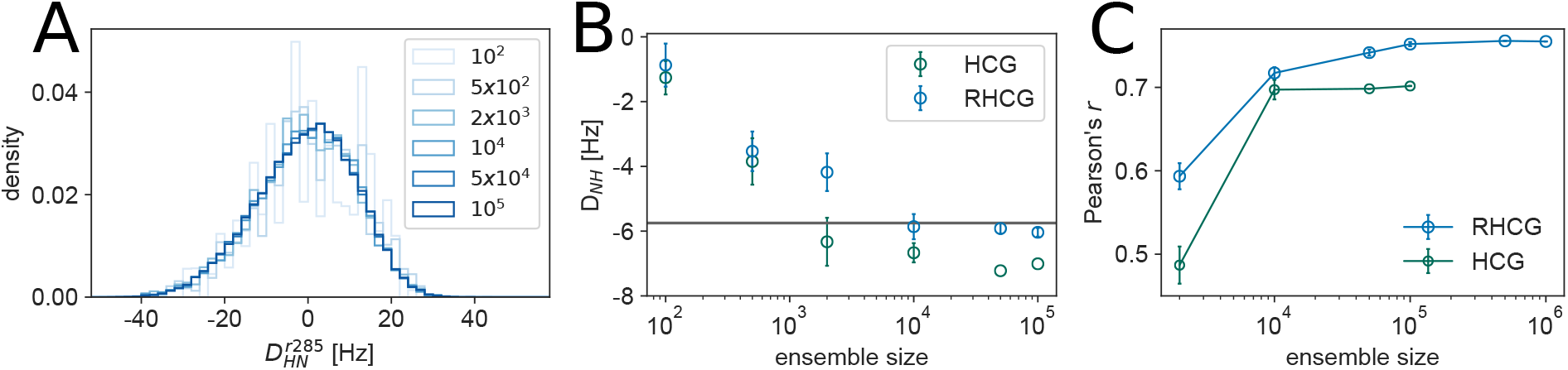
Large ensembles are required to capture NMR RDC measurements. (A) Distribution of ^1^H-^15^N RDC values for L285 in RHCG ensembles of different size, as calculated by PALES^61^ without rescaling. (B) Average ^1^H-^15^N RDC for L285 in dependence of the ensemble size for HCG (dark green) and RHCG (blue). Error bars indicate ±SEM. (C) Ensemble-size dependence of Pearson *r* correlation coefficient between tau K18 ^1^H-^15^N RDC measurements^27^ and calculations from RHCG (blue) and HCG (green), respectively.

The ensemble-size dependence is similar for RHCG and HCG, even if the RHCG ensemble consistently performs somewhat better than the HCG ensemble (Figures 1D, 2B,C and S7).

### The RHCG Ensemble Captures the Extension of tau K18 in Solution

The RHCG ensemble also captures the size and shape of tau K18 in solution as probed by SAXS measurements (Figure 1D). The mean scattering profiles calculated from our tau K18 models agree well with the experimental scattering profiles (Figure 1D), taking possible unspecific aggregation in the low *q* regime into account. The computed root-mean-square radius of gyration of approximately 39 Å coincides with the experimentally determined *R_G_* of 38 ±3 Å.^18^ The RHCG ensemble (〈*R_h_*〉 = 34 Å) is also consistent with the hydrodynamic radius *R_h_* 34±6 Å, as reported by dynamic light scattering (DLS).^71^ *R_h_* was computed from the RHCG ensemble using an empirical approach.^72,73^ Moreover, our RHCG ensemble agrees quite well with NMR paramagnetic relaxation enhancement (PRE) measurements (Figure S8), which were also not used in the generation of our ensembles. Spin-label dynamics were modelled with a rotamer-library approach^74^ and the overall shapes of the experimental profiles measured for four different spin-labels^71^ were captured without any refinement.^46^ In particular, the widths of the valleys with reduced PREs close to the spin label are in line with the experimental profiles and thus the compaction of the chain on the scale of 20-30 amino acids is captured well. The good agreement with SAXS, dynamic light scattering, and NMR measurements suggests that the RHCG ensemble captures the global conformational properties of tau K18 in solution quite well without further refinement.

### Structure of tau K18 as Assessed by Single-Molecule FRET

Comparison to single-molecule FRET experiments suggests that our RHCG models are somewhat too extended (Figure 1E), with longer C*α*-C*α* distances in the RHCG ensemble than those extracted from the FRET experiments. ^45^ However, in a BioEn calculation we found that already a small correction of the chain weights suffices to match the mean distance deduced from FRET perfectly (Figure S2D), with a Kullback-Leibler divergence of *S*_KL_ ≈ 0.2 corresponding to a change of the underlying MD simulation potential energy function of *S*_KL_*k*_B_*T* = ∫ *d***x***p*^(opt)^ (**x**) [*U*^(opt)^(**x**) – *U*(**x**)] ≈ 0.5 kJ/mol on average.^39^ Conversely, this sensitivity also highlights the intricacies of the free energy landscape of disordered proteins, where subtle shifts in the energetics result in appreciable changes in conformation. ^75^

We explored possible effects of the fluorescent dyes by generating RHCG models with dyes attached. For these models, we calculated the mean FRET efficiency and compared it directly to the experimental measurement (Figure S2C). We found that an even smaller force field correction of 0.35 kJ/mol on average^39^ would be sufficient to achieve full consistency of the ensemble means (Figure S2D). Overall, reweighting according to the FRET data does not appreciably affect the agreement with SAXS. Reweighting according to the FRET data changes the *R*_G_ from 39.4 Å (RHCG) to 37.4 Å (RHCG*), and with explicit dye models from 40.1 Å (RHCG+dyes) to 39.1 Å (RHCG+dyes*), respectively.

The scaling exponent of 0.57 inferred from the SAW-*ν* model^66^ is close to the value of an excluded-volume chain. The tau K18 segment is thus more extended than most moderately charged disordered IDPs. ^21^ Interestingly, the transfer efficiency and average distance between the Cys residues of tau K18 from single-molecule FRET are virtually independent of salt concentration (Figure S2C), indicating that the rather pronounced expansion of this segment is not caused by charge repulsion. The FRET experiments are thus in line with our modeling which highlights that residual structure rather than charge-charge interactions shape the ensemble of tau K18 and its overall extension and shape.

### Aggregation-Prone Extended Structures Feature Prominently in the Solution Ensemble of tau K18

Interestingly, a small but significant fraction of our atomic resolution models feature conformations of the two aggregation-prone hexapeptide motifs^10^ as seen in the high-resolution structures of tau fibrils. ^76,77^ Chain growth thus captures biologically important structural features. For the first hexapeptide motif ^275^VQIINK^280^, we found that about 9% of the models are within 1 Å C_*α*_ RMSD of a tau fragment fibril structure (PDB: 5V5B^77^) (Figure 3A,C). A similar fraction of the tau K18 population has local structures matching that of a fibril from a corticobasal degeneration (CBD) patient sample^78^ (PDB: 6TJO). The fraction of our ensemble that closely matches the experimental structures (Figure 3B and Figure S9) is clearly larger than what would be expected for a random six amino acid segment. For the second hexapeptide motif ^306^VQIVYK^311^, we also found that about 8% of the models are within 1.0 Å *C_α_* RMSD of the X-ray structure (PDB: 2ON9 ^76^) (Figure 3B,D), about 2.5 times more than what would be expected for random hexapeptide segments. We found similar consistency for the second hexapeptide motif with the structures of tau fibrils (Figure S9), as formed in Alzheimer’s disease (PDB: 5O30,^79^ 5O3T,^79^ 6HRE,^80^ 6HRF^80^), CBD (6TJO,^78^ 6VI3^81^), Pick’s disease (6GX5^82^) and chronic traumatic encephalopathy (6NWP^83^). Experiments on tau K18 in solution suggest that these motifs should be partially in extended conformations, consistent with our ensemble. ^16,27^

**Figure 3:**
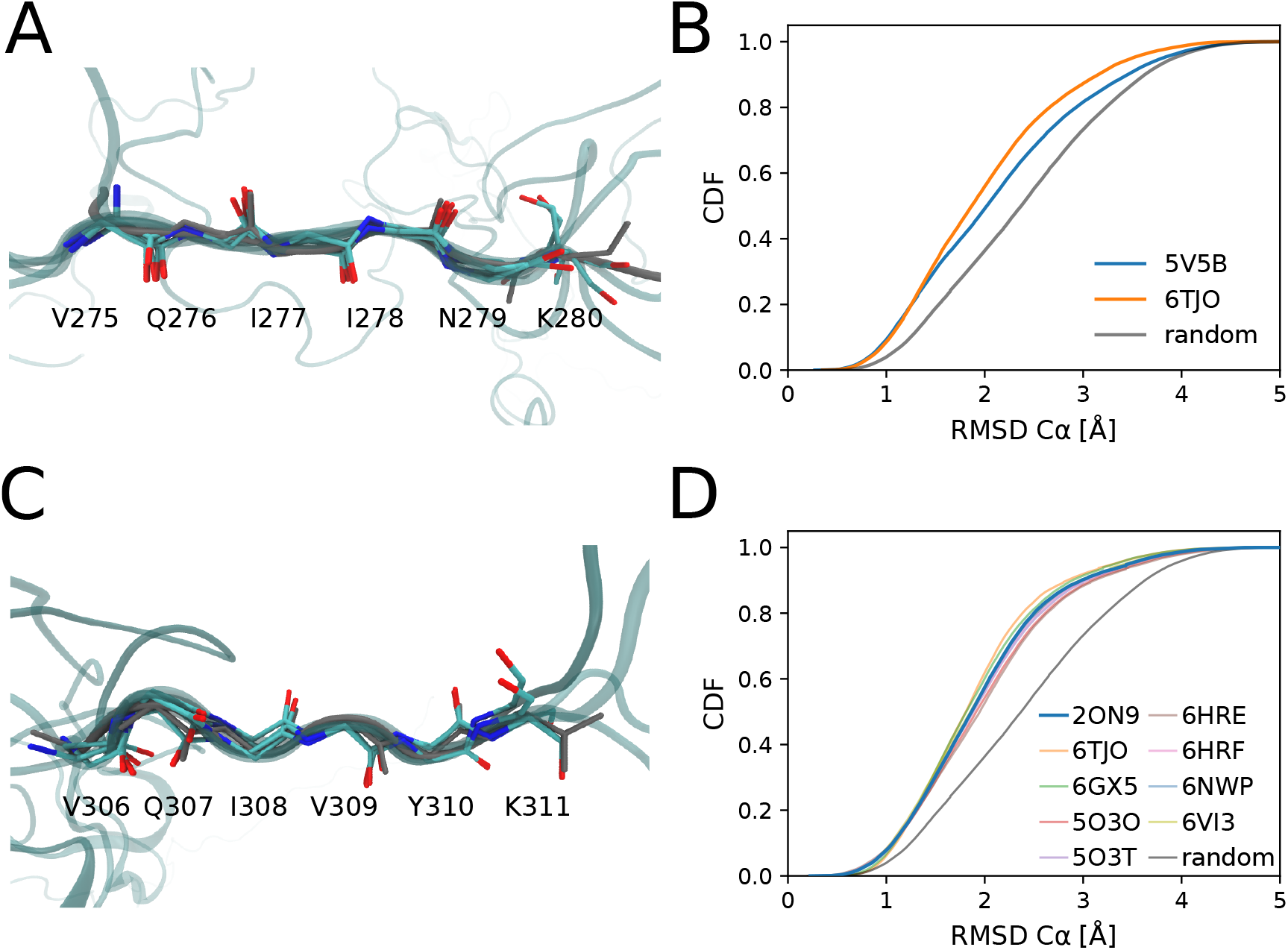
Atomic-resolution RHCG ensembles feature the extended conformations seen in high-resolution structures of tau fibrils. (A,B) ^275^VQIINK^280^ and (C,D) ^306^VQIVYK^311^ hexapeptide motifs are compared to their experimental structures in tau fibrils. (A,C) Five RHCG structures (C_*α*_ RMSD < 0.5 Å from RHCG are superimposed on experimental structures (gray, PDB: 5V5B and 2ON9). (B) Cumulative distribution of RMSD to experimental structure. For reference, gray line show the distributions obtained for the RMSD between 50000 randomly chosen six amino-acid segments in our model ensembles and the motifs in 5V5B. (D) Cumulative distribution of RMSD to experimental structure. For reference, the gray line shows the distribution of the RMSD between randomly chosen six amino-acid segments and the hexapeptide motif in the fibril (PDB: 2ON9).

### The Solution Ensemble Contains the Functional Conformations of tau in Complex with Microtubules

We found that a considerable fraction of WT tau K18 adopts locally compact turn-like structures (Figure 4A-C). Similar turn-like structures have been resolved by NMR NOESY experiments probing the conformations of microtubule-bound tau,^54^ with an O(300)-N(303) distance below 4 Å in 18 out of the 20 structures in the NMR ensemble (PDB: 2MZ7; see Figure 4C). In the WT RHCG ensemble, 15 % of structures of the ^300^VPGGG^304^ segment are within 1 Å C_*α*_ RMSD of the closest representative of the NMR ensemble (Figure 4A). This provides a strong indication that tau samples the turn-like structures of the microtubule-bound form also free in solution.

**Figure 4:**
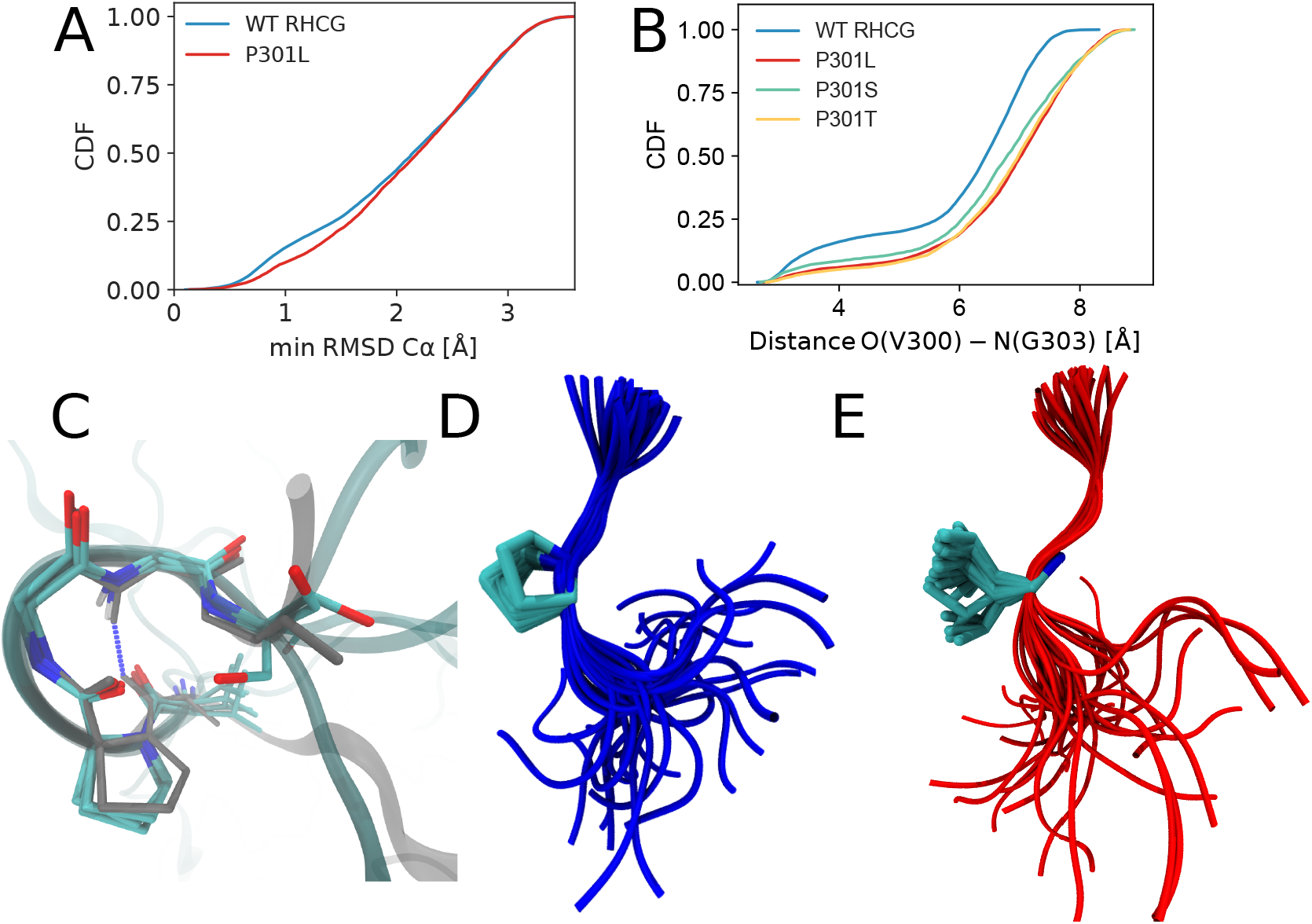
Tau P301 mutations favor more extended local structures. (A) Cumulative distributions of the minimum C_*α*_ RMSD of ^300^VPGGG^304^ to the closest representative of the NMR structural ensemble.^54^ Results are shown for the RHCG ensembles of WT tau K18 and for the HCG ensemble of the P301L variant. (B) Cumulative distributions of O(V300)-N(G303) distances for WT tau K18 from RHCG compared to P301L, P301S and P301T tau K18 variants from HCG. (C) Five representative structures of ^300^VPGGG^304^ from RHCG (oxygen: red; nitrogen: blue; carbon: cyan; C_*α*_ RMSD < 0.5 Å) are superimposed on a representative of the NMR structural ensemble (gray sticks, PDB: 2MZ7, structure 17). Tubes indicate the amino-acid backbone. The O(300)-N(303) hydrogen bond is indicated by the blue dashed line. (D) Representative local structures of WT tau K18. (E) Representative local structures of the P301L variant. In D and E, the structures were aligned on residues 300 and 301. Tubes indicate the backbone. Side-chain heavy atoms and amide nitrogen and C_*α*_ of residue 301 are shown as licorice.

### RHCG Captures the Effect of Mutations Towards Aggregation-Prone Structures

The PG motifs at the end of each repeat favor turn structures. ^84^ We expect that mutations of the prolines shift the local structure away from turns. To test the effect of mutations at the 301 position, we considered the mutations P301L, P301S and P301T. The P301L mutation has been shown to strongly promote tau aggregation. ^9,10^

According to chemical shift mapping, P301 mutations do not affect the global structure of tau.^11^ However, in our hierarchical modeling the P301L, P301S and P301T variants consistently form more extended structures than WT, both in ensembles of full-length tau K18 (Figure 4B-E) and in fragment MD simulations (Figure S10). This loss of turn-like structures is indicated by a more than twofold reduction in the fraction of O-N distances <4 Å between V300 and G303. The shift from turns to extended structures rationalizes the enhanced aggregation propensity of tau P301L in vitro^10,12^ because extended structures predominate in fibrils. Locally more extended structures in the mutant proteins facilitate intermolecular contacts between tau chains and subsequent assembly and aggregation via intermolecular *β*-sheets. The shift to extended structures seen here also explains why P301L tau binds less strongly to microtubules.^11,85^ In a population-shift mechanism, P301L, P301S and P301T mutations thus appear to decrease the fraction of tau with locally compact turn structures, which are competent to bind to microtubules, and to increase the fraction of aggregation-prone extended structures (Figures 5 and S11). The combination of these two effects may render P301 mutations deleterious both with respect to a loss in function and an increased tendency to form disease-associated fibrils.

**Figure 5:**
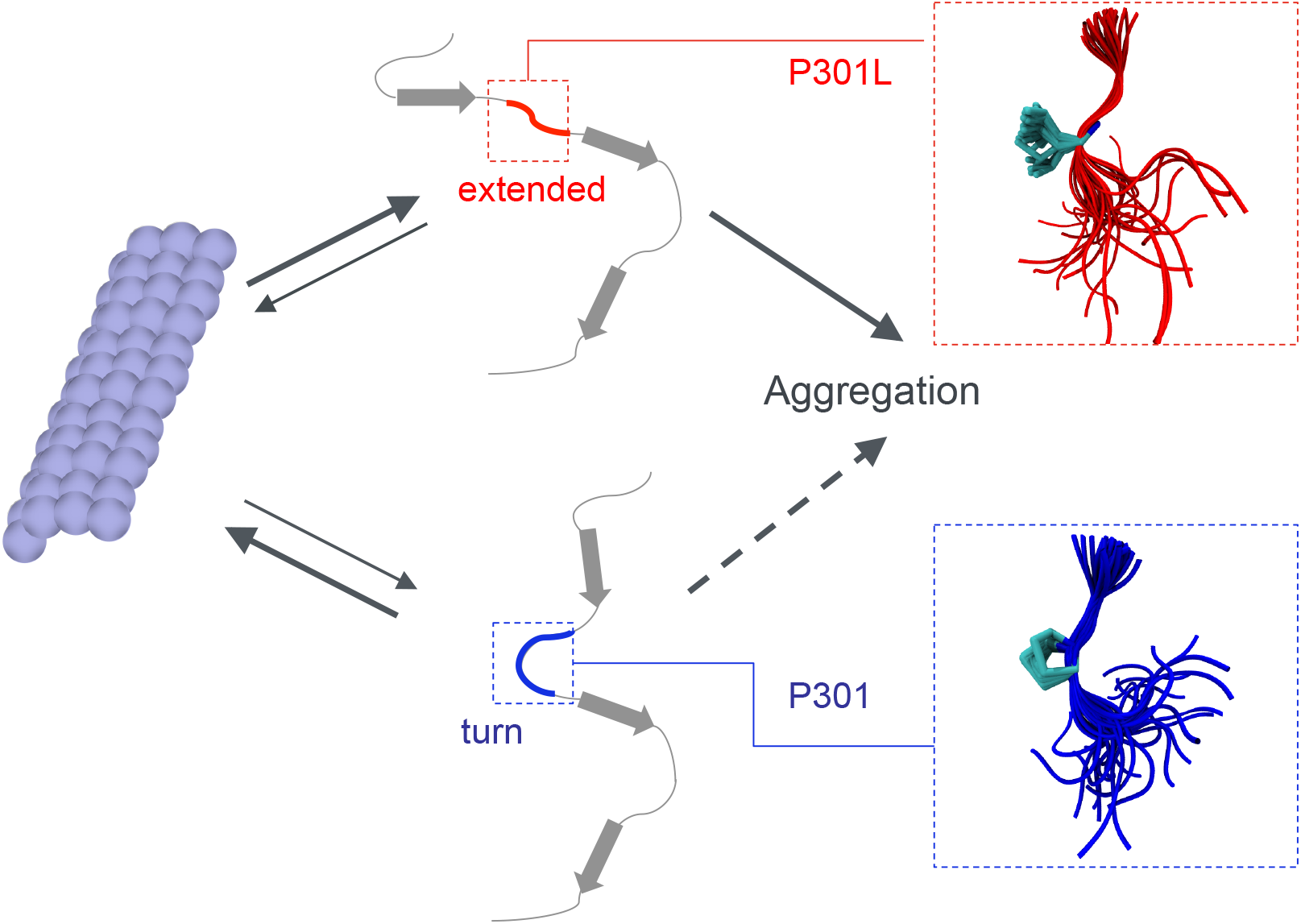
P301 mutations shift the balance from functional to aggregation-prone conformations. Turn conformations (bottom) are required for functional microtubule binding (left), whereas extended conformations (top) are associated with aggregation and the formation of pathogenic fibrils (right). In the wild-type ensemble (P301; bottom), turn-like structures predominate. By contrast, extended structures are significantly populated in the mutant ensemble (P301L; top). The zoom-ins on the right show representative backbone traces around amino acid 301 as tubes.

## Conclusions

We showed that reweighted hierarchical chain growth captures both the local and the global structures of tau K18. Locally, NMR C_*α*_ chemical shifts were reproduced within the expected uncertainties without any fitting. The agreement was improved further with only a gentle Bayesian ensemble refinement against NMR chemical shift data. Globally, the tau K18 chains assembled in this way reproduced SAXS, FRET and NMR RDC measurements, and thus captured the overall shape, dimension and changes in orientation. The global structure of tau K18 thus emerged from its local structure.

Fragment assembly and coil models have proved remarkably successful in the modeling of disordered proteins. ^24-26,28,36,52,55,70^ The quality of the ensemble models can be improved even further by integrating experimental data.^26,35^ However, if the summed squared error *χ*^2^ of the models grows roughly linearly with chain length, e.g., because of systematic errors in the force field used to generate the fragment models, the relative weights of the assembled chains in a refined ensemble will vary widely. As a result, the overlap between the ensemble of assembled chains and the true ensemble becomes exponentially small and ensemble refinement becomes increasingly inefficient.

Reweighted hierarchical chain growth is an importance sampling procedure designed to address this problem by producing evenly weighted ensembles. By applying a bias already during chain assembly, we ensure that the assembled chains have near-uniform weights in the final ensemble. By using hierarchical chain growth^52^ and correcting for any bias in the assembly process in a formally rigorous manner using a form of Bayesian ensemble refinement, BioEn, ^39^ we ensure further that the final ensemble is well defined and independent of arbitrary choices in the assembly process, such as the strength of the bias in fragment selection or the direction of chain growth.

The tau K18 ensembles obtained by reweighted hierarchical chain growth revealed how deleterious mutations shift the balance from protein function to disease. In modeling the effect of mutations, we took advantage of a chemically informed description^70,86-93^ of the disordered tau protein. We found that already free in solution, the microtubule-interacting regions of tau K18 populate local structures as observed in the microtubule-bound state by NMR. Also consistent with conformational selection, we found that a comparable fraction of free tau K18 chains exhibits local structures as observed in pathogenic tau fibrils. We could further show that the disease-associated mutations P301L, P301S and P301T shift the balance away from the microtubule-bound local turn structures toward the fibril-associated extended structures (Figure 5). Such shifts can have dramatic effects on the kinetics of aggregation^94^ by lowering the barrier to nucleation. Indeed, a shift to extended structures was recently reported to be associated with fibril formation in tau condensates. ^95^ The emergence of global structure from local structure thus extends beyond chain shape, dimension and orientation to the competition between tau’s role as microtubule-bound regulator of cellular transport and as fibril-forming driver of neuropathologies.

## Supporting Information Available

Structural and topological analysis of tau K18 ensembles, Comparison to C_*α*_ chemical shifts, construction of fragment pool, modeling of P301 mutations, global BioEn reweighting, analysis of turn structures detected by RDCs, comparison to PRE experiments, comparison to NMR structure of microtubule bound ^301^PGGG^304^, details on the sample preparation and setup of single-molecule FRET experiments and further comparison of single-molecule FRET experiments and ensembles from RHCG, analysis of the structural consequences of P301 mutations, sequence of tau K18 and positions of spin labels and FRET dyes.

## Acknowledgement

We acknowledge financial support from the German Research Foundation (CRC902: Molecular Principles of RNA Based Regulation), the Max Planck Society, and the Swiss National Science Foundation. M.Z. was supported by the European Research Council (ERC) under the EU Horizon 2020 research and innovation program (grant agreement No. 787679). L.S.S. thanks ReALity (Resilience, Adaptation and Longevity), M^3^ODEL (Mainz Instiute of Multiscale Modeling) and Forschungsinitiative des Landes Rheinland-Pfalz for their support and Prof. Marek Cieplak and Dr. Peter Virnau for insightful discussions. M.S. was supported by the FWF Schrödinger fellowship J4332-B28. We thank Daniel Nettels for providing data analysis tools for single-molecule spectroscopy.

## References

(1) Kadavath, H.; Hofele, R. V.; Biernat, J.; Kumar, S.; Tepper, K.; Urlaub, H.; Mandelkow, E.; Zweckstetter, M. Tau Stabilizes Microtubules by Binding at the Interface Between Tubulin Heterodimers. Proc. Natl. Acad. Sci. U. S. A. 2015, 112, 7501–6.

(2) Wang, Y.; Mandelkow, E. Tau in Physiology and Pathology. Nat. Rev. Neurosci. 2016, 17, 22–35.

(3) Hernandez-Vega, A.; Braun, M.; Scharrel, L.; Jahnel, M.; Wegmann, S.; Hyman, B. T.; Alberti, S.; Diez, S.; Hyman, A. A. Local Nucleation of Microtubule Bundles through Tubulin Concentration into a Condensed Tau Phase. Cell Reports 2017, 20, 2304–2312.

(4) Ambadipudi, S.; Biernat, J.; Riedel, D.; Mandelkow, E.; Zweckstetter, M. Liquid-Liquid Phase Separation of the Microtubule-Binding Repeats of the Alzheimer-Related Protein Tau. Nat. Commun. 2017,

(5) Zhang, X.; Lin, Y.; Eschmann, N. A.; Zhou, H.; Rauch, J. N.; Hernandez, I.; Guzman, E.; Kosik, K. S.; Han, S. RNA Stores Tau Reversibly in Complex Coacervates. PLOS Biol. 2017, 15, e2002183.

(6) Ukmar-Godec, T.; Wegmann, S.; Zweckstetter, M. Biomolecular Condensation of the Microtubule-Associated Protein Tau. Semin. Cell Dev. Biol. 2020, 99, 202–214, SI: Protein-Protein Interactions in Health and Disease.

(7) Wegmann, S.; Eftekharzadeh, B.; Tepper, K.; Zoltowska, K. M.; Bennett, R. E.; Dujardin, S.; Laskowski, P. R.; MacKenzie, D.; Kamath, T.; Commins, C.; Vanderburg, C.; Roe, A. D.; Fan, Z.; Molliex, A. M.; Hernandez-Vega, A.; Muller, D.; Hyman, A. A.; Mandelkow, E.; Taylor, J. P.; Hyman, B. T. Tau Protein Liquid-Liquid Phase Separation Can Initiate Tau Aggregation. EMBO J. 2018, 37, e98049.

(8) Lathuilière, A.; Valdés, P.; Papin, S.; Cacquevel, M.; Maclachlan, C.; Knott, G. W.; Muhs, A.; Paganetti, P.; Schneider, B. L. Motifs in the Tau Protein That Control Binding to Microtubules and Aggregation Determine Pathological Effects. Sci. Rep. 2017, 7, 13556.

(9) Strang, K. H.; Croft, C. L.; Sorrentino, Z. A.; Chakrabarty, P.; Golde, T. E.; Giasson, B. I. Distinct Differences in Prion-Like Seeding and Aggregation Between Tau Protein Variants Provide Mechanistic Insights Into Tauopathies. J. Biol. Chem. 2018, 293, 2408–2421.

(10) Von Bergen, M.; Barghorn, S.; Li, L.; Marx, A.; Biernat, J.; Mandelkow, E. M.; Mandelkow, E. Mutations of Tau Protein in Frontotemporal Dementia Promote Aggregation of Paired Helical Filaments by Enhancing Local *β*-Structure. J. Biol. Chem. 2001,

(11) Fischer, D.; Mukrasch, M. D.; von Bergen, M.; Klos-Witkowska, A.; Biernat, J.; Griesinger, C.; Mandelkow, E.; Zweckstetter, M. Structural and Microtubule Binding Properties of Tau Mutants of Frontotemporal Dementias. Biochemistry 2007, 46, 2574–2582.

(12) Chen, D.; Drombosky, K. W.; Hou, Z.; Sari, L.; Kashmer, O. M.; Ryder, B. D.; Perez, V. A.; Woodard, D. R.; Lin, M. M.; Diamond, M. I.; Joachimiak, L. A. Tau Local Structure Shields an Amyloid-Forming Motif and Controls Aggregation Propensity. Nat. Commun. 2019, 10, 2493.

(13) Bonomi, M.; Heller, G. T.; Camilloni, C.; Vendruscolo, M. Principles of Protein Structural Ensemble Determination. Curr. Opin. Struct. Biol. 2017, 42, 106–116.

(14) Schwalbe, M.; Ozenne, V.; Bibow, S.; Jaremko, M.; Jaremko, L.; Gajda, M.; Jensen, M. R.; Biernat, J.; Becker, S.; Mandelkow, E.; Zweckstetter, M.; Blackledge, M. Predictive Atomic Resolution Descriptions of Intrinsically Disordered hTau40 and *α*-Synuclein in Solution from NMR and Small Angle Scattering. Structure 2014, 22, 238–249.

(15) Meier, S.; Blackledge, M.; Grzesiek, S. Conformational Distributions of Unfolded Polypeptides From Novel NMR Techniques. J. Chem. Phys. 2008, 128, 052204.

(16) Mukrasch, M. D.; Bibow, S.; Korukottu, J.; Jeganathan, S.; Biernat, J.; Griesinger, C.; Mandelkow, E.; Zweckstetter, M. Structural Polymorphism of 441-Residue Tau at Single Residue Resolution. PLoS Biol. 2009, 7, 1–1.

(17) Jensen, M. R.; Salmon, L.; Nodet, G.; Blackledge, M. Defining Conformational Ensembles of Intrinsically Disordered and Partially Folded Proteins Directly From Chemical Shifts. J. Am. Chem. Soc. 2010, 132, 1270–1272.

(18) Mylonas, E.; Hascher, A.; Bernadó, P.; Blackledge, M.; Mandelkow, E.; Svergun, D. I. Domain Conformation of Tau Protein Studied by Solution Small-Angle X-Ray Scattering. Biochemistry 2008, 47 10345–10353.

(19) Kalinin, S.; Peulen, T.; Sindbert, S.; Rothwell, P. J.; Berger, S.; Restle, T.; Goody, R. S.; Gohlke, H.; Seidel, C. A. M. A Toolkit and Benchmark Study for FRET-Restrained High-Precision Structural Modeling. Nat. Methods 2012, 9, 1218–1225.

(20) Shammas, S. L.; Garcia, G. A.; Kumar, S.; Kjaergaard, M.; Horrocks, M. H.; Shivji, N.; Mandelkow, E.; Knowles, T. P.; Mandelkow, E.; Klenerman, D. A Mechanistic Model of Tau Amyloid Aggregation Based on Direct Observation of Oligomers. Nat. Commun. 2015, 6, 7025.

(21) Schuler, B.; Soranno, A.; Hofmann, H.; Nettels, D. Single-Molecule FRET Spectroscopy and the Polymer Physics of Unfolded and Intrinsically Disordered Proteins. Annu. Rev. Biophys. 2016, 45, 207–231.

(22) Borgia, A.; Borgia, M. B.; Bugge, K.; Kissling, V. M.; Heidarsson, P. O.; Fernandes, C. B.; Sottini, A.; Soranno, A.; Buholzer, K. J.; Nettels, D.; Kragelund, B. B.; Best, R. B.; Schuler, B. Extreme Disorder in an Ultrahigh-Affinity Protein Complex. Nature 2018, 555, 61–66.

(23) Brucale, M.; Schuler, B.; Samorì, B. Single-Molecule Studies of Intrinsically Disordered Proteins. Chem. Rev. 2014, 114, 3281–3317.

(24) Fiebig, K. M.; Schwalbe, H.; Buck, M.; Smith, L. J.; Dobson, C. M. Toward a Description of the Conformations of Denatured States of Proteins. Comparison of a Random Coil Model With NMR Measurements. J. Phys. Chem. 1996, 100, 2661–2666.

(25) Schwalbe, H.; Fiebig, K. M.; Buck, M.; Jones, J. A.; Grimshaw, S. B.; Spencer, A.; Glaser, S. J.; Smith, L. J.; Dobson, C. M. Structural and Dynamical Properties of a Denatured Protein. Heteronuclear 3D NMR Experiments and Theoretical Simulations of Lysozyme in 8 M Urea. Biochemistry 1997, 36, 8977–8991.

(26) Bernadó, P.; Bertoncini, C. W.; Griesinger, C.; Zweckstetter, M.; Blackledge, M. Defining Long-Range Order and Local Disorder in Native *α*-Synuclein Using Residual Dipolar Couplings. J. Am. Chem. Soc. 2005, 127, 17968–17969.

(27) Mukrasch, M. D.; Markwick, P.; Biernat, J.; von Bergen, M.; Bernadó, P.; Griesinger, C.; Mandelkow, E.; Zweckstetter, M.; Blackledge, M. Highly Populated Turn Conformations in Natively Unfolded Tau Protein Identified from Residual Dipolar Couplings and Molecular Simulation. J. Am. Chem. Soc. 2007, 129, 5235–5243.

(28) Fisher, C. K.; Huang, A.; Stultz, C. M. Modeling Intrinsically Disordered Proteins With Bayesian Statistics. J. Am. Chem. Soc. 2010, 132, 14919–14927.

(29) Mantsyzov, A. B.; Maltsev, A. S.; Ying, J.; Shen, Y.; Hummer, G.; Bax, A. A Maximum Entropy Approach to the Study of Residue-Specific Backbone Angle Distributions in *α*-Synuclein, an Intrinsically Disordered Protein. Protein Sci. 2014, 23, 1275–1290.

(30) Brotzakis, Z. F.; Lindstedt, P. R.; Taylor, R.; Bernardes, G. J. L.; Vendruscolo, M. A Structural Ensemble of a Tau-Microtubule Complex Reveals Regulatory Tau Phosphorylation and Acetylation Mechanisms. bioRxiv 2020,

(31) Aznauryan, M.; Delgado, L.; Soranno, A.; Nettels, D.; Huang, J. R.; Labhardt, A. M.; Grzesiek, S.; Schuler, B. Comprehensive Structural and Dynamical View of an Unfolded Protein From the Combination of Single-Molecule FRET, NMR, and SAXS. Proc. Natl. Acad. Sci. U. S. A. 2016, 113, E5389–98.

(32) Borgia, A.; Zheng, W.; Buholzer, K.; Borgia, M. B.; Schüler, A.; Hofmann, H.; Soranno, A.; Nettels, D.; Gast, K.; Grishaev, A.; Best, R. B.; Schuler, B. Consistent View of Polypeptide Chain Expansion in Chemical Denaturants From Multiple Experimental Methods. J. Am. Chem. Soc. 2016, 138, 11714–11726.

(33) Escobedo, A.; Topal, B.; Kunze, M. B. A.; Aranda, J.; Chiesa, G.; Mungianu, D.; Bernardo-Seisdedos, G.; Eftekharzadeh, B.; Gairí, M.; Pierattelli, R.; Felli, I. C.; Diercks, T.; Millet, O.; García, J.; Orozco, M.; Crehuet, R.; Lindorff-Larsen, K.; Salvatella, X. Side Chain to Main Chain Hydrogen Bonds Stabilize a Polyglutamine Helix in a Transcription Factor. Nat. Commun. 2019, 10, 2034.

(34) Rezaei-Ghaleh, N.; Parigi, G.; Soranno, A.; Holla, A.; Becker, S.; Schuler, B.; Luchinat, C.; Zweckstetter, M. Local and Global Dynamics in Intrinsically Disordered Synuclein. Angew. Chemie Int. Ed. 2018, 57, 15262–15266.

(35) Gomes, G.-N. W.; Krzeminski, M.; Namini, A.; Martin, E. W.; Mittag, T.; Head-Gordon, T.; Forman-Kay, J. D.; Gradinaru, C. C. Conformational Ensembles of an Intrinsically Disordered Protein Consistent With NMR, SAXS, and Single-Molecule FRET. J. Am. Chem. Soc. 2020, 142, 15697–15710.

(36) Krzeminski, M.; Marsh, J. A.; Neale, C.; Choy, W.-Y.; Forman-Kay, J. D. Characterization of Disordered Proteins With ENSEMBLE. Bioinformatics 2013, 29, 398–399.

(37) Rieping, W.; Habeck, M.; Nilges, M. Inferential Structure Determination. Science 2005, 309, 303–306.

(38) Różycki, B.; Kim, Y. C.; Hummer, G. SAXS Ensemble Refinement of ESCRT-III CHMP3 Conformational Transitions. Structure 2011, 19, 109–116.

(39) Hummer, G.; Köfinger, J. Bayesian Ensemble Refinement by Replica Simulations and Reweighting. J. Chem. Phys. 2015, 143, 243150.

(40) Köfinger, J.; Stelzl, L. S.; Reuter, K.; Allande, C.; Reichel, K.; Hummer, G. Efficient Ensemble Refinement by Reweighting. J. Chem. Theory Comput. 2019, 15, 3390–3401.

(41) Bonomi, M.; Camilloni, C.; Cavalli, A.; Vendruscolo, M. Metainference: A Bayesian Inference Method for Heterogeneous Systems. Sci. Adv. 2016, 2.

(42) Boomsma, W.; Tian, P. F.; Frellsen, J.; Ferkinghoff-Borg, J.; Hamelryck, T.; Lindorff-Larsen, K.; Vendruscolo, M. Equilibrium Simulations of Proteins Using Molecular Fragment Replacement and NMR Chemical Shifts. Proc. Natl. Acad. Sci. U.S.A. 2014, 111, 13852–13857.

(43) Bottaro, S.; Bengtsen, T.; Lindorff-Larsen, K. In Structural Bioinformatics: Methods and Protocols; Gáspári, Z., Ed.; Springer US: New York, NY, 2020; pp 219–240.

(44) Best, R. B. Computational and Theoretical Advances in Studies of Intrinsically Disordered Proteins. Curr. Opin. Struct. Biol. 2017, 42, 147–154.

(45) Holmstrom, E. D.; Holla, A.; Zheng, W.; Nettels, D.; Best, R. B.; Schuler, B. Accurate Transfer Efficiencies, Distance Distributions, and Ensembles of Unfolded and Intrinsically Disordered Proteins From Single-Molecule FRET. Methods Enzymol. 2018, 611, 287–325.

(46) Reichel, K.; Stelzl, L. S.; Köfinger, J.; Hummer, G. Precision DEER Distances From Spin-Label Ensemble Refinement. J. Phys. Chem. Lett. 2018, 9, 5748–5752.

(47) Salvi, N.; Abyzov, A.; Blackledge, M. Multi-Timescale Dynamics in Intrinsically Disordered Proteins From NMR Relaxation and Molecular Simulation. J. Phys. Chem. Lett. 2016, 7, 2483–2489.

(48) Löhr, T.; Jussupow, A.; Camilloni, C. Metadynamic Metainference: Convergence Towards Force Field Independent Structural Ensembles of a Disordered Peptide. J. Chem. Phys. 2017, 146, 165102.

(49) Larsen, A. H.; Wang, Y.; Bottaro, S.; Grudinin, S.; Arleth, L.; Lindorff-Larsen, K. Combining Molecular Dynamics Simulations With Small-Angle X-Ray and Neutron Scattering Data to Study Multi-Domain Proteins in Solution. PLOS Comput. Biol. 2020, 16, e1007870.

(50) Feldman, H. J.; Hogue, C. W. A Fast Method to Sample Real Protein Conformational Space. Proteins: Structure, Function, and Bioinformatics 2000, 39, 112–131.

(51) Lettieri, S.; Mamonov, A. B.; Zuckerman, D. M. Extending Fragment-Based Free Energy Calculations With Library Monte Carlo Simulation: Annealing in Interaction Space. J. Comput. Chem. 2011, 32, 1135–1143.

(52) Pietrek, L. M.; Stelzl, L. S.; Hummer, G. Hierarchical Ensembles of Intrinsically Disordered Proteins at Atomic Resolution in Molecular Dynamics Simulations. J. Chem. Theory Comput. 2020, 16, 725–737.

(53) Lincoff, J.; Haghighatlari, M.; Krzeminski, M.; Teixeira, J. M.; Gomes, G.-N. W.; Gradinaru, C. C.; Forman-Kay, J. D.; Head-Gordon, T. Extended Experimental Inferential Structure Determination Method in Determining the Structural Ensembles of Disordered Protein States. Commun. Chem. 2020, 3, 1–12.

(54) Kadavath, H.; Jaremko, M.; Jaremko, Ł.; Biernat, J.; Mandelkow, E.; Zweckstetter, M. Folding of the Tau Protein on Microtubules. Angew. Chemie Int. Ed. 2015, 54, 10347–10351.

(55) Mamonov, A. B.; Bhatt, D.; Cashman, D. J.; Ding, Y.; Zuckerman, D. M. General Library-Based Monte Carlo Technique Enables Equilibrium Sampling of Semi-Atomistic Protein Models. J. Phys. Chem. B 2009, 113, 10891–10904.

(56) Shen, Y.; Bax, A. SPARTA+: A Modest Improvement in the Empirical NMR Chemical Shift Predicition by Means of an Artifical Neural Network. J. Biomol. NMR 2011, 48, 13–22.

(57) Nielsen, J. T.; Mulder, F. A. POTENCI: Prediction of Temperature, Neighbor and pH-corrected Chemical Shifts for Intrinsically Disordered Proteins. J. Biomol. NMR 2018, 70, 141–165.

(58) Vögeli, B.; Ying, J.; Grishaev, A.; Bax, A. Limits on Variations in Protein Backbone Dynamics From Precise Measurements of Scalar Couplings. J. Am. Chem. Soc. 2007, 129, 9377–9385.

(59) McGibbon, R. T.; Beauchamp, K. A.; Harrigan, M. P.; Klein, C.; Swails, J. M.; Hernández, C. X.; Schwantes, C. R.; Wang, L.-P.; Lane, T. J.; Pande, V. S. MDTraj: A Modern Open Library for the Analysis of Molecular Dynamics Trajectories. Biophys. J. 2015, 109, 1528–1532.

(60) Zweckstetter, M.; Bax, A. Prediction of Sterically Induced Alignment in a Dilute Liquid Crystalline Phase: Aid to Protein Structure Determination by NMR. J. Am. Chem. Soc. 2000, 122, 3791–3792.

(61) Zweckstetter, M. NMR: Prediction of Molecular Alignment From Structure Using the PALES Software. Nat. Protoc. 2008, 3, 679–690.

(62) Bax, A.; Kontaxis, G.; Tjandra, N. Dipolar Couplings in Macromolecular Structure Determination. Methods Enzymol. 2001, 339, 127–74.

(63) Schneidman-Duhovny, D.; Hammel, M.; Tainer, J. A.; Sali, A. FoXS, FoXSDock and MultiFoXS: Single-State and Multi-State Structural Modeling of Proteins and Their Complexes Based on SAXS Profiles. Nucleic Acids Res. 2016, 44, W424–W429.

(64) Michaud-Agrawal, N.; Denning, E. J.; Woolf, T. B.; Beckstein, O. MDAnalysis: A Toolkit for the Analysis of Molecular Dynamics Simulations. J. Comput. Chem. 2011, 32, 2319–2327.

(65) Gowers, R. J.; Linke, M.; Barnoud, J.; Reddy, T. J. E.; Melo, M. N.; Seyler, S. L.; Dotson, D. L.; Domański, J.; Buchoux, S.; Kenney, I. M.; Beckstein, O. MDAnalysis: A Python Package for the Rapid Analysis of Molecular Dynamics Simulations. Proceedings of the 15th Python in Science Conference. 2016; pp 98–105.

(66) Zheng, W.; Zerze, G. H.; Borgia, A.; Mittal, J.; Schuler, B.; Best, R. B. Inferring Properties of Disordered Chains From FRET Transfer Efficiencies. J. Chem. Phys. 2018, 148, 123329.

(67) Grotz, K. K.; Nuuesch, M. F.; Holmstrom, E. D.; Heinz, M.; Stelzl, L. S.; Schuler, B.; Hummer, G. Dispersion Correction Alleviates Dye Stacking of Single-Stranded DNA and RNA in Simulations of Single-Molecule Fluorescence Experiments. J. Phys. Chem. B 2018, 122, 11626–11639.

(68) Franke, D.; Petoukhov, M. V.; Konarev, P. V.; Panjkovich, A.; Tuukkanen, A.; Mertens, H. D. T.; Kikhney, A. G.; Hajizadeh, N. R.; Franklin, J. M.; Jeffries, C. M.; Svergun, D. I. ATSAS 2.8: A Comprehensive Data Analysis Suite for Small-Angle Scattering From Macromolecular Solutions. J. Appl. Crystallogr. 2017, 50, 1212–1225.

(69) Robustelli, P.; Piana, S.; Shaw, D. E. Developing a Molecular Dynamics Force Field for Both Folded and Disordered Protein States. Proc. Natl. Acad. Sci. 2018, 115, E4758–E4766.

(70) Estaña, A.; Sibille, N.; Delaforge, E.; Vaisset, M.; Cortés, J.; Bernadó, P. Realistic Ensemble Models of Intrinsically Disordered Proteins Using a Structure-Encoding Coil Database. Structure 2019, 27, 381–391.

(71) Akoury, E.; Mukrasch, M. D.; Biernat, J.; Tepper, K.; Ozenne, V.; Mandelkow, E.; Blackledge, M.; Zweckstetter, M. Remodeling of the Conformational Ensemble of the Repeat Domain of Tau by an Aggregation Enhancer. Protein Sci. 2016, 25, 1010–1020.

(72) Nygaard, M.; Kragelund, B. B.; Papaleo, E.; Lindorff-Larsen, K. An Efficient Method for Estimating the Hydrodynamic Radius of Disordered Protein Conformations. Biophys. J. 2017, 113, 550–557.

(73) Ahmed, M. C.; Crehuet, R.; Lindorff-Larsen, K. Computing, Analyzing and Comparing the Radius of Gyration and Hydrodynamic Radius in Conformational Ensembles of Intrinsically Disordered Proteins. bioRxiv 2019,

(74) Tesei, G.; Martins, J. M.; Kunze, M. B. A.; Wang, Y.; Crehuet, R.; Lindorff-Larsen, K. DEER-PREdict: Software for Efficient Calculation of Spin-Labeling EPR and NMR Data From Conformational Ensembles. PLOS Comput. Biol. 2021, 17, 1–18.

(75) Yeh, I. C.; Hummer, G. Peptide Loop-Closure Kinetics From Microsecond Molecular Dynamics Simulations in Explicit Solvent. J. Am. Chem. Soc. 2002, 124, 6563–6568.

(76) Sawaya, M. R.; Sambashivan, S.; Nelson, R.; Ivanova, M. I.; Sievers, S. A.; Apostol, M. I.; Thompson, M. J.; Balbirnie, M.; Wiltzius, J. J. W.; McFarlane, H. T.; Madsen, A. Ø.; Riekel, C.; Eisenberg, D. Atomic Structures of Amyloid Cross-*β* Spines Reveal Varied Steric Zippers. Nature 2007, 447, 453–457.

(77) Seidler, P. M.; Boyer, D. R.; Rodriguez, J. A.; Sawaya, M. R.; Cascio, D.; Murray, K.; Gonen, T.; Eisenberg, D. S. Structure-Based Inhibitors of Tau Aggregation. Nat. Chem. 2018, 10, 170–176.

(78) Zhang, W.; Tarutani, A.; Newell, K. L.; Murzin, A. G.; Matsubara, T.; Falcon, B.; Vidal, R.; Garringer, H. J.; Shi, Y.; Ikeuchi, T.; Murayama, S.; Ghetti, B.; Hasegawa, M.; Goedert, M.; Scheres, S. H. W. Novel Tau Filament Fold in Corticobasal Degeneration. Nature 2020, 580, 283–287.

(79) Fitzpatrick, A. W. P.; Falcon, B.; He, S.; Murzin, A. G.; Murshudov, G.; Garringer, H. J.; Crowther, R. A.; Ghetti, B.; Goedert, M.; Scheres, S. H. W. Cryo-Em Structures of Tau Filaments From Alzheimer’s Disease. Nature 2017, 547, 185–190.

(80) Falcon, B.; Zhang, W.; Schweighauser, M.; Murzin, A. G.; Vidal, R.; Garringer, H. J.; Ghetti, B.; Scheres, S. H. W.; Goedert, M. Tau Filaments From Multiple Cases of Sporadic and Inherited Alzheimer’s Disease Adopt a Common Fold. Acta Neuropathol. 2018, 136, 699–708.

(81) Arakhamia, T.; Lee, C. E.; Carlomagno, Y.; Duong, D. M.; Kundinger, S. R.; Wang, K.; Williams, D.; DeTure, M.; Dickson, D. W.; Cook, C. N.; Seyfried, N. T.; Petrucelli, L.; Fitzpatrick, A. W. Posttranslational Modifications Mediate the Structural Diversity of Tauopathy Strains. Cell 2020, 180, 633–644.e12.

(82) Falcon, B.; Zhang, W.; Murzin, A. G.; Murshudov, G.; Garringer, H. J.; Vidal, R.; Crowther, R. A.; Ghetti, B.; Scheres, S. H. W.; Goedert, M. Structures of Filaments From Pick’s Disease Reveal a Novel Tau Protein Fold. Nature 2018, 561, 137–140.

(83) Falcon, B.; Zivanov, J.; Zhang, W.; Murzin, A. G.; Garringer, H. J.; Vidal, R.; Crowther, R. A.; Newell, K. L.; Ghetti, B.; Goedert, M.; Scheres, S. H. W. Novel Tau Filament Fold in Chronic Traumatic Encephalopathy Encloses Hydrophobic Molecules. Nature 2019, 568, 420–423.

(84) Mukrasch, M. D.; Biernat, J.; von Bergen, M.; Griesinger, C.; Mandelkow, E.; Zweckstetter, M. Sites of Tau Important for Aggregation Populate *β*-Structure and Bind to Microtubules and Polyanions. J. Biol. Chem. 2005, 280, 24978–24986.

(85) Hong, M.; Zhukareva, V.; Vogelsberg-Ragaglia, V.; Wszolek, Z.; Reed, L.; Miller, B. I.; Geschwind, D. H.; Bird, T. D.; McKeel, D.; Goate, A.; Morris, J. C.; Wilhelmsen, K. C.; Schellenberg, G. D.; Trojanowski, J. Q.; Lee, V. M.-Y. Mutation-Specific Functional Impairments in Distinct Tau Isoforms of Hereditary FTDP-17. Science 1998, 282, 1914–1917.

(86) Mioduszewski, Ł.; Różycki, B.; Cieplak, M. Pseudo-Improper-Dihedral Model for Intrinsically Disordered Proteins. J. Chem. Theory Comput. 2020, 16, 4726–4733.

(87) Vitalis, A.; Pappu, R. V. ABSINTH: A New Continuum Solvation Model for Simulations of Polypeptides in Aqueous Solutions. J. Comput. Chem. 2009, 30, 673–699.

(88) Baul, U.; Chakraborty, D.; Mugnai, M. L.; Straub, J. E.; Thirumalai, D. Sequence Effects on Size, Shape, and Structural Heterogeneity in Intrinsically Disordered Proteins. J. Phys. Chem. B 2019, 123, 3462–3474.

(89) Wu, H.; Wolynes, P. G.; Papoian, G. A. AWSEM-IDP: A Coarse-Grained Force Field for Intrinsically Disordered Proteins. J. Phys. Chem. B 2018, 122, 11115–11125.

(90) Zhao, Y.; Cortes-Huerto, R.; Kremer, K.; Rudzinski, J. F. Investigating the Conformational Ensembles of Intrinsically Disordered Proteins With a Simple Physics-Based Model. J. Phys. Chem. B 2020, 124, 4097–4113.

(91) Benayad, Z.; von Bülow, S.; Stelzl, L. S.; Hummer, G. Simulation of FUS Protein Condensates With an Adapted Coarse-Grained Model. J. Chem. Theory Comput. 2020, 525–537.

(92) Gruijs da Silva, L. A.; Simonetti, F.; Hutten, S.; Riemenschneider, H.; Sternburg, E. L.; Pietrek, L. M.; Gebel, J.; Doetsch, V.; Edbauer, D.; Hummer, G.; Stelzl, L. S.; Dormann, D. Disease-Linked TDP-43 Hyperphosphorylation Suppresses TDP-43 Condensation and Aggregation. bioRxiv 2021, 2021.04.30.442163.

(93) Farr, S. E.; Woods, E. J.; Joseph, J. A.; Garaizar, A.; Collepardo-Guevara, R. Nucleosome Plasticity is a Critical Element of Chromatin Liquid–Liquid Phase Separation and Multivalent Nucleosome Interactions. Nat. Commun. 2021, 12, 2883.

(94) Chakraborty, D.; Straub, J. E.; Thirumalai, D. Differences in the Free Energies Between the Excited States of A*β*40 and A*β*42 Monomers Encode Their Aggregation Propensities. Proc. Natl. Acad. Sci. U. S. A. 2020, 117, 19926–19937.

(95) Wen, J.; Hong, L.; Krainer, G.; Yao, Q.-Q.; Knowles, T. P. J.; Wu, S.; Perrett, S. Conformational Expansion of Tau in Condensates Promotes Irreversible Aggregation. J. Am. Chem. Soc. 2021, 143, 13056–13064.

