## Supporting Information for "Global Structure of the Intrinsically Disordered Protein Tau Emerges from its Local Structure"

∇*Rocasolano Institute for Physical Chemistry, CSIC, Serrano 119, 28006 Madrid, Spain*

### Contents

**Supporting Text** **S-2**

Structural and Topological Analysis of tau K18 Ensembles . . . . . . . . . . . . . S-2

Comparison of Structural Ensembles to NMR $C_\alpha$ Chemical Shifts . . . . . . . . . . S-3

Construction of Biased Fragment Pool . . . . . . . . . . . . . . . . . . . . . . . . S-3

Mutations at P301 Position . . . . . . . . . . . . . . . . . . . . . . . . . . . . . S-4

Global BioEn Reweighting . . . . . . . . . . . . . . . . . . . . . . . . . . . . . S-4

Analysis of Local Turn Structures Detected by RDCs . . . . . . . . . . . . . . . . S-5

Comparison of Structural Ensembles to PRE Experiments . . . . . . . . . . . . . S-5

Structure of $^{301}$PGGG$^{304}$ Motif of Wild-Type and P301L tau K18 . . . . . . . . . . S-7

Further Analysis of the Structural Consequences of P301 Mutations . . . . . . . . S-8

Labeling of tau K18 with Fluorescent Dyes . . . . . . . . . . . . . . . . . . . . . S-8

Single-Molecule Fluorescence Spectroscopy . . . . . . . . . . . . . . . . . . . . . S-9

Comparison of Structural Ensembles to FRET Experiments . . . . . . . . . . . . S-10

**References** **S-12**

**Supporting Table S1** **S-15**

**Supporting Figures S1–S11** **S-16**

### Supporting Text

#### Structural and Topological Analysis of tau K18 Ensembles

We found that both HCG and RHCG ensembles are highly diverse. The mean pairwise $\mathrm{RMSD}_{S_i,S_j}$ values of ≈26 Å (Fig. S3) are large and comparable to the 27 Å RMSD found previously for HCG ensembles of $\alpha$-synuclein.[S1]

We also tested whether the tau K18 structures featured knots. To this end we applied the Koniaris-Muthukumar-Taylor (KMT) reduction algorithm,[S2,S3] which progressively shrinks the polymer chain by averaging positions of neighboring monomers while disallowing chain crossings. If a straight line connects the termini after the final iteration, the chain is considered unknotted. We did not find any knots in ensembles of 50000 structures obtained by HCG and RHCG. Indeed, for short self-avoiding random walks or short and weakly interacting polymers few knots are expected.[S4,S5]

#### Comparison of Structural Ensembles to NMR $C_\alpha$ Chemical Shifts

NMR chemical shifts are accurate reporters of average local structure. The magnitude of the secondary $C_\alpha$ chemical shifts calculated from the tau K18 ensemble obtained by reweighted hierarchical chain growth (RHCG) are small in general as are the experimental shifts (Figure S4B). In key stretches, e.g., for residues S285 to V300, the ensemble from RHCG reproduces the overall trend from experiment. However, even though the error in the chemical shift prediction is relatively large at $\approx$1 ppm and our calculated shifts are thus well within the uncertainty at each position, we noticed that our models quite consistently produced slightly larger shifts than those measured by NMR with a mean unsigned error of 0.27 ppm. This trend indicates a slight overabundance of simulation conformers in the $\alpha$-helical minimum of the Ramachandran map at the expense of locally extended structures. To improve the representation of the local structure in our initial ensemble from HCG, as probed by the NMR chemical shifts (Figure S4A, with a mean unsigned error of 0.35 ppm), we applied a reweighting scheme.

#### Construction of Biased Fragment Pool

For RHCG, we adapted the hierarchical chain growth (HCG) algorithm described in ref S1. The peptide fragment conformations collected from replica-exchange molecular dynamics (REMD), as described in the main text, entered the fragment library with a probability weight factor. The subsequent assembly of the fragments from this biased pool proceeded as in HCG.

The fragment weights were determined according to the NMR chemical-shift data. For each fragment position except the last, $1 \leq n < 43$, we performed separate Bayesian inference of ensemble (BioEn) refinements[S6,S7] against the respective $C_\alpha$ chemical shift data measured for the central residues in each fragment (i.e., excluding residues 1 and 5). For the last fragment, $n = 43$, we used uniform weights. With a confidence parameter of $\theta = 10$, the REMD ensemble was only gently biased, as indicated by typically small Kullback-Leibler (KL) divergences $S_{\mathrm{KL}}^{\mathrm{BioEn}}$ of $0.004 \pm 2 \times 10^{-5}$ across the 42 fragments. The maximum $S_{\mathrm{KL}}^{\mathrm{BioEn}}$ was 0.019.

#### Mutations at P301 Position

We studied the effect of disease-associated mutations at the P301 positions. In the original division of tau K18 into overlapping pentamer fragments, P301 appeared at the second position. To minimize possible end effects, we repeated the chain growth with a second set of fragments, in which P301 occupied the central position of its pentamer fragment. Shifting residue P301 to the central position of the fragment changed the residue overlap with the preceding and following fragments to 3 and 1 residues, respectively. Using the fragment library with centered 301 position, we assembled 10 000 wild-type, P301L, P301S and P301T models with HCG.

#### Global BioEn Reweighting

The bias in the assembly of the RHCG ensembles was removed in a final BioEn refinement. RHCG shifted the population of residual structures more towards extended conformers with less positive chemical shifts compared to HCG (Figure S4A,B). As a result, the Pearson $r$ correlation coefficient improved from $r = 0.28$ for HCG to $r = 0.41$ for RHCG (Figure S4C,D). Note that with expected uncertainties of about 1 ppm, any further improvement would likely result in overfitting.

We set $\theta = 5$ in the global BioEn reweighting according to the L-curve analysis in Figure S1A. For $\theta = 5$, we found $S_{\text{KL}}^{\text{BioEn}} < 1$ $S_{\text{KL}}^{\text{bias}} \ll 1$. The fragments produced by the REMD simulations are thus already quite consistent with the NMR chemical-shift data, and the bias in chain growth produced tau K18 structures of nearly uniform weight in the final RHCG ensemble (Figure S1B).

#### Analysis of Local Turn Structures Detected by RDCs

Previous NMR residual dipolar coupling[S8] experiments have identified four turns in tau K18: $^{252}$DLKN$^{255}$, $^{283}$DLSN$^{286}$, $^{314}$DLSK$^{317}$ and $^{345}$DFKD$^{348}$. For the first two turn motifs, RHCG captured the trend in RDCs well, as shown in Figure 1 in the main text. Local distances also suggest a tendency to form turns (Figure S6). By contrast, the control segment V275-I278 is more extended (Figure S6) and the RDCs do not show a distinct pattern (Figure 1A in the main text). For the third turn motif at position 345-348, RHCG captured the drop in the RDCs but not the subsequent rise. $C_\alpha(n) - C^{'}(n+3)$ distances also suggest relatively extended structures and not a clear turn. These distances are consistent with one sharp change in direction and a less well defined second change in direction. The fourth turn motif is broadly captured. $C_\alpha(n) - C^{'}(n+3)$ distances show a significant compact population indicative of a turn and the RDCs broadly agree between experiment and our models RHCG.

#### Comparison of Structural Ensembles to PRE Experiments

Paramagnetic relaxation enhancement (PRE) measurements constitute another source of long-range distance information from NMR experiments. PREs were computed using the PREpredict[S9] Python library (https://github.com/KULL-Centre/DEERpredict). PREpredict uses a rotamer library approach to model the different possible conformation of the spin label and attach the spin label to each structure in an ensemble of protein structures. The dynamics of the spin label is described by a model-free approach. The relaxation enhancement due to the electron spin on the paramagnetic spin label is then computed from the Solomon-Bloembergen equation. PREs for the backbone amides were computed for spin labels positioned at residues I260, C291, C322 and I354 (Table S1) to compare to the PRE measurements in aqueous solution by Akoury et al.[S10] For the calculation, we set $\tau_c = 4$ ns and $\tau_t = 0.2$ ns. To meet the experimental conditions we set T = 278 K, the total INEPT-delays in the pulse sequence were $\approx 10$ ms and $\omega = 2\pi \times 800$ MHz. $R_{2,\mathrm{dia}}$ was set based on NMR spin-relaxation experiments. The values were in the range of $5 \pm 1$ s$^{-1}$. The PREpredict analysis was run with 20 different tau ensembles of size $k = 50\,000$, which makes a total of 1 million models. PREpredict returns the weighted ensemble average, using the respective weights from global BioEn reweighting. For each label we calculated the median and the interquartile range for each residue from the distribution of the 20 averages. From the distribution of the 20 averages we calculated the median for each label and the interquartile range.

Overall, we find quite good agreement of the PREs predicted for the tau ensembles from RHCG with the experimental profile (Figure S8). Our tau ensembles feature models that reproduce the local compaction of tau K18, indicated by the width of the valley of the PRE profiles. However, according to the PREs predicted for our tau ensembles we over- and underestimate some distances compared to the PREs reported by the experiment. E.g., we observe discrepancies for the label at residue 291 with the label being too close to the flanking residues 270-290 and too far away from residues 335-360 (compare Figure S8B). A possible source of such mismatches may be that in PRE experiments the relaxation rates are enhanced due to the dipolar coupling between an unpaired electron, e.g., from a paramagnetic label, and a nucleus. The interaction is dependent on the $\langle \mathrm{r}^{-6} \rangle$ average distance between the dipolar coupled species.[S11] This allows for the measurement of sparsely populated states in the fast exchange regime. I.e., one or a few instances with the unpaired electron and the proton very close to each other increase the relaxation rate and conversely decrease the PRE intensity ratio significantly. In other words, if in a 50 000 tau subensemble only one S-(1-

oxyl-2,2,5,5-tetramethyl-2,5-dihydro-1H-pyrrol-3-yl)methyl methanesulfonothioate (MTSL) rotamer is mapped onto a tau conformation such that a residue of interest is very close to the label with r≈0, the distance for this rotamer-backbone combination significantly affects the $\langle r^{-6} \rangle$ average. Consequently, the predicted PRE profile for this residue is decreased. One way to address this issue is to perform a reweighting of the initial rotamer weights in our ensemble using,[S12] e.g., by BioEn,[S6] to deal with outliers and thus further improve the agreement with the experimental PRE profiles.

#### Structure of $^{301}$PGGG$^{304}$ Motif of Wild-Type and P301L tau K18

Encouragingly, the compact conformational state of the $^{301}$PGGG$^{304}$ motif we have identified is consistent with previous NMR investigations of wild-type tau bound to microtubules.[S13] The agreement does not depend whether P301 is the second position or the in the central position of the fragment used for hierarchical chain growth, as shown in Figure S11. Microtubule-bound tau adopts turn-like and compact conformations in this region and some of these local conformations show a high degree of similarity with the local structure in our full-length tau models. Specifically, locally compact turn structures for $^{300}$VPGGG$^{304}$ are found in our ensemble and in the NMR structure of microtubule-associated tau.[S13] For the five residue stretch $^{300}$VPGGG$^{304}$ about 15 % of the ensemble is within 1.0 Å minimum $C_\alpha$ RMSD to one of the 20 NMR structures (Figure 4C and S11). The NMR structures of microtubule-bound tau make a sharp turn at position 304 and show some tertiary contacts, which are not present in the ensemble from hierarchical chain growth. The agreement between our modeling and the NMR structure is highly localized, but indicates nonetheless that structural features detected in experiments emerge naturally in our modeling.

The ensemble for the P301L variant features fewer conformations similar to the microtubule-bound NMR structure[S13] than wild-type tau K18. About 10 % of the P301L ensemble are within 1 Å $C_\alpha$ RMSD of one of the microtubule-bound NMR structures, compared to 15 % for WT (Figure S11). This shift is consistent with the distance analysis of the O-N atoms of V300 and G303 (Figure 4B in the main text and Figure S10). Note that growing models of mutant tau K18 with P301L at the central position of the respective fragment does not change the overall conclusion, and would suggest an even larger effect of the P301L mutant, with about 4 % within 1.0 Å minimum $C_\alpha$ RMSD to one of the 20 NMR structures.

#### Further Analysis of the Structural Consequences of P301 Mutations

We found that P301 mutants are locally more extended than wild-type tau K18. We consistently found these differences in our full-length models and on the level of atomistic MD simulations of individual fragments. For P301 mutants we found a decreased population of these locally compact structures and the mutants adopt locally a more extended structure. The differences between wild-type tau K18 and P301 mutants are apparent at the level of peptide fragments irrespective whether P301 is at the second or the central position (Figure S10A and B, respectively) of the pentamer fragments.

#### Labeling of tau K18 with Fluorescent Dyes

For the single-molecule experiments, tau K18 was labeled with Alexa Fluor 488 and CF660R at C291 and C322 (Table S1). Freshly reduced tau K18 was purified by RP-HPLC using a Reprosil Gold C18 column (Dr. Maisch GmbH, Germany) with a water-acetonitrile (AcN) gradient (Solvent A: 0.1 % TFA, Solvent B: AcN; gradient: 5-50 % B in 30 min). The protein was labeled at the natural Cys residues with the donor dye Alexa Fluor 488 maleimide (Molecular Probes, Eugene, USA) in 6 M guanidinium chloride (GdmCl), 100 mM sodium phosphate, pH 7.3, overnight at 4°C at a molar ratio of dye to protein of 0.7:1. After stopping the reaction with DTT (1,4-dithiotreitol), singly labeled material was purified using a gradient of 24.5-28.5 % B on the same column. The purified material was further labeled with the acceptor dye CF660R maleimide (Biotium, Fremont, USA) overnight at 4°C using a molar ratio of dye to protein of 5:1. Unreacted dye was quenched with DTT. Purification of donor-acceptor labeled protein was achieved using a 25-30% gradient of B. The mass of donor/acceptor-labeled tau K18 was confirmed by ESI-MS.

#### Single-Molecule Fluorescence Spectroscopy

For the single-molecule experiments, tau K18 labeled with Alexa Fluor 488 and CF660R was diluted to 100 pM in 50 mM sodium phosphate buffer, pH 6.8, 1 mM DTT, 0.001 % Tween 20 or 20 mM HEPES, 5 mM KCl, 10 mM $MgCl_2$, pH 7.4, 1 mM DTT, 0.001% Tween 20. The experiments were performed at 295 K on a MicroTime 200 confocal single-molecule instrument (PicoQuant, Berlin, Germany) using chambered polymer coverslips (µ-Slide, ibidi, Gräfelfing, Germany). Excitation light was provided by a pulsed diode laser (LDH-D-C-485, Pico-Quant, Berlin, Germany) at a power of 55 µW at a pulse repetition rate of 20 MHz for the donor dye, and a white-light continuum source (SuperK EXTREME EXW12, NKT photonics, Brøndby, Denmark) filtered with a z582/15 band-pass filter (Chroma, Brattleboro, USA) at a power of 120 µW at 20 MHz for the acceptor dye. Both power values were measured at the back aperture of the objective (UplanApo 60x/1.20 W, Olympus, Tokyo, Japan). Fluorescence light passed through a 100-nm pinhole and was then split by polarization and color (595 DCXR, Chroma, Brattleboro, USA). After passing additional filters (ET 525/50, Chroma, Brattleboro, USA, for donor fluorescence and Edge Basic LP635, Semrock, Rochester, USA, for acceptor fluorescence), the emitted light was detected with four avalanche photodiodes (SPCM-AQR-15, PerkinElmer Optoelectronics, Vaudreuil, QC, Canada). Arrival times of detected photons were recorded with the timing resolution set to 16 ps (HydraHarp 400, PicoQuant, Berlin, Germany).

Pulsed interleaved excitation[S14] was used to determine all relevant correction factors for these measurements from a set of samples labeled with Alexa Fluor 488 and CF660R as described by Lee et al. and Holmstrom et al.[S15,S16] Time bins of 1 ms with more than 50 photons after donor or acceptor excitation were selected as bursts. The corresponding photon counts were then corrected using the correction factors for differences in quantum yield of the dyes, detection efficiencies, spectral crosstalk, and direct excitation of the acceptor dye by the donor excitation laser.[S16] For each burst, transfer efficiency and stoichiometry ratio were calculated, and the corresponding mean values for donor-acceptor labeled molecules were determined by 2D-Gaussian fits to stoichiometry-versus-transfer efficiency plots. From the resulting mean transfer efficiencies, the average Cys-to-Cys distances were inferred with the SAW-$\nu$ model for the distance distribution[S17] and corrected for the contribution of the dye linkers by length rescaling, assuming that each dye and linker is equivalent to 4.5 amino acid residues.[S16] The dye pair Alexa Fluor 488/CF660R was used because of its low Förster radius of 4.6 nm (calculated from the donor emission and acceptor absorption spectra of the corresponding free dyes), which makes it well suited for probing the short sequence separation between the Cys residues in tau K18. The Förster radius was adjusted for the refractive index of the solution according to Förster's theory.[S18] All single-molecule data analysis was conducted using Fretica, a custom Wolfram Symbolic Transfer Protocol (WSTP) add-on for Mathematica (Wolfram), available at https://schuler.bioc.uzh.ch/programs/.

#### Comparison of Structural Ensembles to FRET Experiments

To compute FRET efficiencies for our structural ensembles of tau K18, we constructed models with fluorescent dyes. In RHCG+dyes, we added the two dyes in the final step of fragment assembly. A chain and a pair of dyes were drawn at random. If no clash was found, the dyes were added to the chain. If a clash was found, a new chain and two new dyes were drawn. As an approximate model for both the acceptor and donor dyes, we used the Alexa Fluor 488 library from Grotz et al.[S19] Chains were drawn according to their weights from a RHCG ensemble of 100,000 independently generated chains. In this way, we generated 50,000 tau structures with dyes.

The C$\alpha$-C$\alpha$ distances for the ensembles grown without or with dyes are very nearly indistinguishable (Figure S2A). However, as expected, the calculated interdye-distance is about 5 Å larger than the C$\alpha$-C$\alpha$ distance, with a root-mean-squared distance of 51 Å

(Figure S2B). Therefore, as a further test, we calculated FRET efficiencies and compared them to the measurements.

Steady-state donor and acceptor fluorescence anisotropies from the single-molecule measurements were below 0.1, indicating sufficiently fast and isotropic averaging of the relative fluorophore orientations for assuming the orientational factor $\kappa$ in Förster theory to be $2/3$.[S20] MD simulations of polymeric biomolecules (IDPs and also single-stranded DNA)[S16,S19,S21] also suggest that an orientational factor of 2/3 is appropriate. Since the distance dynamics in disordered proteins on the length scale of tau K18 are much slower than the fluorescence lifetime of the donor, we assume static averaging and calculate mean transfer efficiencies from $\langle E \rangle = \int E(r)P(r)dr$, where $E(r)$ is the transfer efficiency at distance $r$, and $P(r)$ is the normalized distance distribution from our ensemble. The mean transfer efficiency is then given by

$$\langle E \rangle = \int_0^\infty P(r) \left[ 1 + \left( \frac{r}{R_0} \right)^6 \right]^{-1} dr, \tag{1}$$

with $R_0 = 46$ Å, the Förster radius for isotropic dye distributions with $\kappa = 2/3$.

The calculated mean FRET efficiency of the RHCG+dye ensemble is somewhat above the measured value (Figure S2C). However, a mild BioEn ensemble reweighting with a Kullback-Leibler divergence $S_{\mathrm{KL}}^{\mathrm{BioEn}} \approx 0.14$ is sufficient to achieve agreement. The mean C$\alpha$-C$\alpha$ distance in the resulting RHCG+dyes* ensemble is 42 Å, close to the 38 Å inferred with the SAW-$\nu$ model from the FRET data. Part of the small remaining discrepancy may be explained by the fact that the Alexa Fluor 488 used at both label sites in the model has a somewhat shorter linker than the CF660R dye at one of the sites in the experiments. The longer linker would require the C$\alpha$-C$\alpha$ distance to be somewhat shorter. Overall, we conclude that the addition of explicit dyes does not significantly alter the conclusions that the RHCG model is somewhat too extended, which can be corrected by a mild ensemble reweighting.

### References


(S1) Pietrek, L. M.; Stelzl, L. S.; Hummer, G. Hierarchical Ensembles of Intrinsically Disordered Proteins at Atomic Resolution in Molecular Dynamics Simulations. *J. Chem. Theory Comput.* **2020**, *16*, 725–737.

(S2) Koniaris, K.; Muthukumar, M. Knottedness in Ring Polymers. *Phys. Rev. Lett.* **1991**, *66*, 2211–2214.

(S3) Taylor, W. R. A Deeply Knotted Protein Structure and How it Might Fold. *Nature* **2000**, *406*, 916–919.

(S4) Virnau, P.; Kantor, Y.; Kardar, M. Knots in Globule and Coil Phases of a Model Polyethylene. *J. Am. Chem. Soc.* **2005**, *127*, 15102–15106.

(S5) Sumners, D. W.; Whittington, S. G. Knots in Self-avoiding Walks. *J. Phys. A. Math. Gen.* **1988**, *21*, 1689–1694.

(S6) Hummer, G.; Köfinger, J. Bayesian Ensemble Refinement by Replica Simulations and Reweighting. *J. Chem. Phys.* **2015**, *143*, 243150.

(S7) Köfinger, J.; Stelzl, L. S.; Reuter, K.; Allande, C.; Reichel, K.; Hummer, G. Efficient Ensemble Refinement by Reweighting. *J. Chem. Theory Comput.* **2019**, *15*, 3390–3401.

(S8) Mukrasch, M. D.; Markwick, P.; Biernat, J.; von Bergen, M.; Bernadó, P.; Griesinger, C.; Mandelkow, E.; Zweckstetter, M.; Blackledge, M. Highly Populated Turn Conformations in Natively Unfolded Tau Protein Identified from Residual Dipolar Couplings and Molecular Simulation. *J. Am. Chem. Soc.* **2007**, *129*, 5235–5243.

(S9) Tesei, G.; Martins, J. M.; Kunze, M. B. A.; Wang, Y.; Crehuet, R.; Lindorff-Larsen, K. DEER-PREdict: Software for Efficient Calculation of Spin-Labeling EPR and NMR Data From Conformational Ensembles. *PLOS Comput. Biol.* **2021**, *17*, 1–18.

(S10) Akoury, E.; Mukrasch, M. D.; Biernat, J.; Tepper, K.; Ozenne, V.; Mandelkow, E.; Blackledge, M.; Zweckstetter, M. Remodeling of the Conformational Ensemble of the Repeat Domain of Tau by an Aggregation Enhancer. *Protein Sci.* **2016**, *25*, 1010–1020.

(S11) Clore, G. M. Exploring Sparsely Populated States of Macromolecules by Diamagnetic and Paramagnetic NMR Relaxation. *Protein Sci.* **2011**, *20*, 229–246.

(S12) Reichel, K.; Stelzl, L. S.; Köfinger, J.; Hummer, G. Precision DEER Distances From Spin-Label Ensemble Refinement. *J. Phys. Chem. Lett.* **2018**, *9*, 5748–5752.

(S13) Kadavath, H.; Jaremko, M.; Jaremko, Ł.; Biernat, J.; Mandelkow, E.; Zweckstetter, M. Folding of the Tau Protein on Microtubules. *Angew. Chemie Int. Ed.* **2015**, *54*, 10347–10351.

(S14) Müller, B. K.; Zaychikov, E.; Bräuchle, C.; Lamb, D. C. Pulsed Interleaved Excitation. *Biophys. J.* **2005**, *89*, 3508–3522.

(S15) Lee, N. K.; Kapanidis, A. N.; Wang, Y.; Michalet, X.; Mukhopadhyay, J.; Ebright, R. H.; Weiss, S. Accurate FRET Measurements Within Single Diffusing Biomolecules Using Alternating-Laser Excitation. *Biophys. J.* **2005**, *88*, 2939–2953.

(S16) Holmstrom, E. D.; Holla, A.; Zheng, W.; Nettels, D.; Best, R. B.; Schuler, B. Accurate Transfer Efficiencies, Distance Distributions, and Ensembles of Unfolded and Intrinsically Disordered Proteins From Single-Molecule FRET. *Methods Enzymol.* **2018**, *611*, 287–325.

(S17) Zheng, W.; Zerze, G. H.; Borgia, A.; Mittal, J.; Schuler, B.; Best, R. B. Inferring Properties of Disordered Chains From FRET Transfer Efficiencies. *J. Chem. Phys.* **2018**, *148*, 123329.

(S18) Förster, T. Zwischenmolekulare Energiewanderung und Fluoreszenz. *Ann. Phys.* **1948**, *6*, 55–75.

(S19) Grotz, K. K.; Nuuesch, M. F.; Holmstrom, E. D.; Heinz, M.; Stelzl, L. S.; Schuler, B.; Hummer, G. Dispersion Correction Alleviates Dye Stacking of Single-Stranded DNA and RNA in Simulations of Single-Molecule Fluorescence Experiments. *J. Phys. Chem. B* **2018**, *122*, 11626–11639.

(S20) Hellenkamp, B.; Schmid, S.; Doroshenko, O.; Opanasyuk, O.; Kuhnemuth, R.; Rezaei Adariani, S.; Ambrose, B.; Aznauryan, M.; Barth, A.; Birkedal, V.; Bowen, M. E.; Chen, H.; Cordes, T.; Eilert, T.; Fijen, C.; Gebhardt, C.; Gotz, M.; Gouridis, G.; Gratton, E.; Ha, T.; Hao, P.; Hanke, C. A.; Hartmann, A.; Hendrix, J.; Hildebrandt, L. L.; Hirschfeld, V.; Hohlbein, J.; Hua, B.; Hubner, C. G.; Kallis, E.; Kapanidis, A. N.; Kim, J. Y.; Krainer, G.; Lamb, D. C.; Lee, N. K.; Lemke, E. A.; Levesque, B.; Levitus, M.; McCann, J. J.; Naredi-Rainer, N.; Nettels, D.; Ngo, T.; Qiu, R.; Robb, N. C.; Rocker, C.; Sanabria, H.; Schlierf, M.; Schroder, T.; Schuler, B.; Seidel, H.; Streit, L.; Thurn, J.; Tinnefeld, P.; Tyagi, S.; Vandenberk, N.; Vera, A. M.; Weninger, K. R.; Wunsch, B.; Yanez-Orozco, I. S.; Michaelis, J.; Seidel, C. A. M.; Craggs, T. D.; Hugel, T. Precision and Accuracy of Single-Molecule FRET Measurements-a Multi-Laboratory Benchmark Study. *Nat. Methods* **2018**, *15*, 669–676.

(S21) Best, R. B.; Merchant, K. A.; Gopich, I. V.; Schuler, B.; Bax, A.; Eaton, W. A. Effect of Flexibility and Cis Residues in Single-Molecule FRET Studies of Polyproline. *Proc. Natl. Acad. Sci. U. S. A.* **2007**, *104*, 18964–18969.

(S22) Zweckstetter, M. NMR: Prediction of Molecular Alignment From Structure Using the PALES Software. *Nat. Protoc.* **2008**, *3*, 679–690.

#### Supporting Table S1

Table S1: Amino-acid sequence of the 129-residue tau K18 region of tau protein. Labeled residues are highlighted in color. The two cysteine residues labeled in both the single-molecule FRET and PRE experiments are highlighted in gold. Two isoleucine residues only labeled in PRE experiments are highlighted in red-violet.

```
244-QTAPVPMPDL  KNVKSKIGST  ENLKHQPGGG  KVQIINKKLD  LSNVQSKCGS
294-KDNIKHVPGG  GSVQIVYKPV  DLSKVTSKCG  SLGNIHHKPG  GGQVEVKSEK
344-LDFKDRVQSK  IGSLDNITHV  PGGGNKKIE
```

### Supporting Figures S1–S11

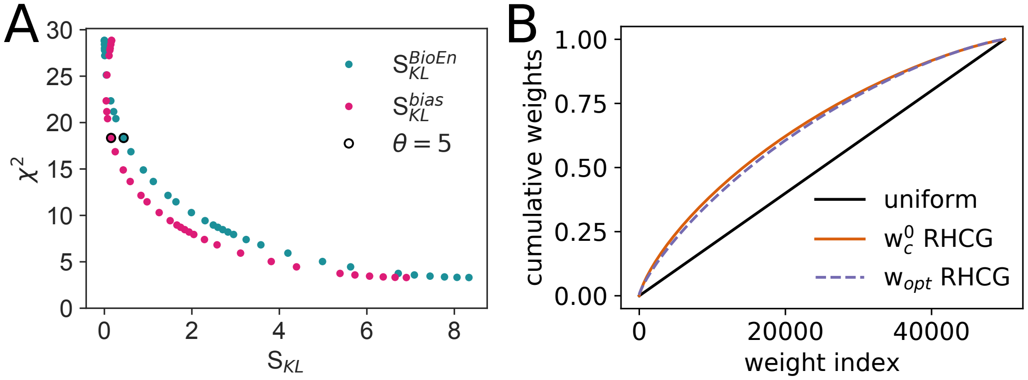


Figure S1: Assessment of the importance sampling approach. (A) L-curve analysis for the global BioEn KL divergence $S_{\mathrm{KL}}^{\mathrm{BioEn}}$ from the initial weights (blue) and the KL divergence $S_{\mathrm{KL}}^{\mathrm{bias}}$ from uniform weights in ideal importance sampling (pink). The black circles indicate the KL divergences for the global BioEn with $\theta = 5$. Results are shown for an ensemble of 50 000 tau K18 chains. The confidence parameter $\theta$ was varied between $10^6$ and 0. $\chi^2$ was calculated according to eq 4 in the main text as the sum over the 109 $C_\alpha$ NMR chemical shifts. Accordingly, the reduced $\chi^2$ is about 0.17 after the global BioEn refinement. (B) Rank ordered cumulative weights plotted against the weight index. The reference chain weights $w_c^0$ before the global BioEn reweighting are shown as solid orange line, the optimal chain weights $w_{opt}$ after global BioEn with $\theta = 5$ as dashed purple line, and a uniform reference as black line.

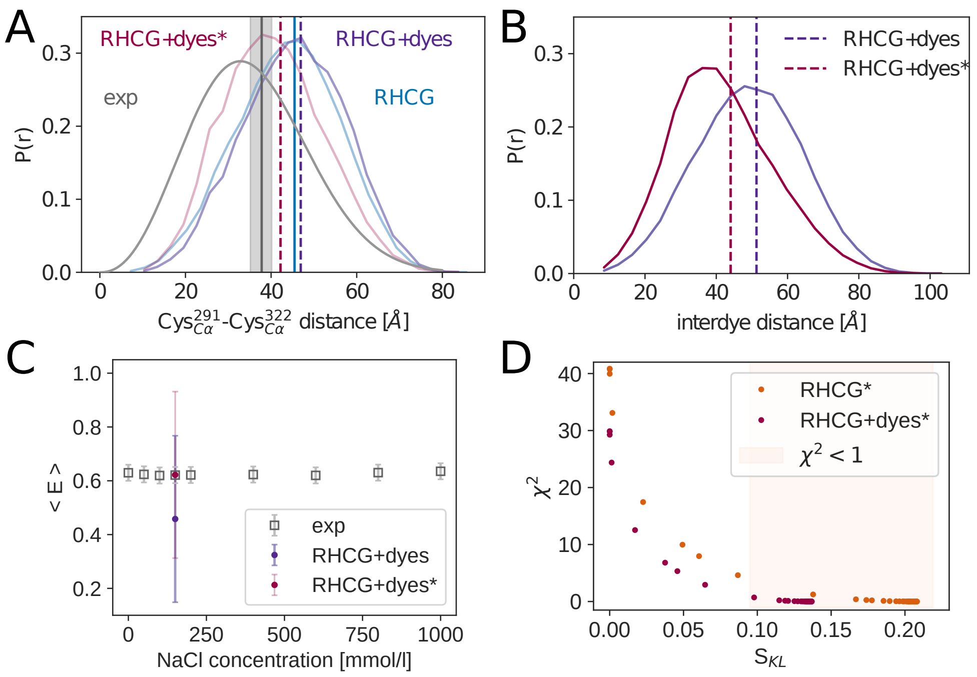


Figure S2: Comparison of Cys$^{291}_{C\alpha}$-Cys$^{322}_{C\alpha}$ distance distributions and FRET efficiencies calculated for RHCG models to single-molecule FRET measurements. (A) Cys$^{291}_{C\alpha}$-Cys$^{322}_{C\alpha}$ distance distribution P(r) as inferred from the experimentally measured FRET efficiencies using the SAW-$\nu$ model[S17] (grey, root-mean-squared average 37.8 Å shown as grey vertical line; grey-shaded area indicates uncertainty based on a 7% error on the Förster radius[S20]), determined for the models from the RHCG ensemble without (blue) and with attached Alexa Fluor 488 dye molecules (purple and dark pink). The RHCG+dye* models were further refined by BioEn reweighting against the experimental FRET efficiency. B) Distribution of interdye distances as calculated for the RHCG+dyes ensemble before and after refinement against the experimental FRET efficiency. The root-mean-squared distance is shown as vertical line. (C) Experimentally measured mean FRET efficiency as a function of NaCl concentration (grey, error bars show the estimated uncertainty of 0.03). For RHCG+dyes(*) models, the FRET efficiency was calculated using the Förster equation (purple and dark magenta, respectively). We show the standard deviation of the distribution of determined efficiencies for RHCG+dyes(*) as error bars to indicate the width of the distribution. (D) L-curves showing the $\chi^2$ values plotted against the $S_{\mathrm{KL}}$ values as a result for the BioEn reweighting against experimental C$\alpha$ root-mean-squared distances of the RHCG ensemble and for the RHCG+dyes ensemble reweighted against the experimental FRET efficiency (orange and dark magenta, respectively). The light salmon shaded area indicates the $S_{\mathrm{KL}}^{\mathrm{BioEn}}$ values for which $\chi^2 < 1$.

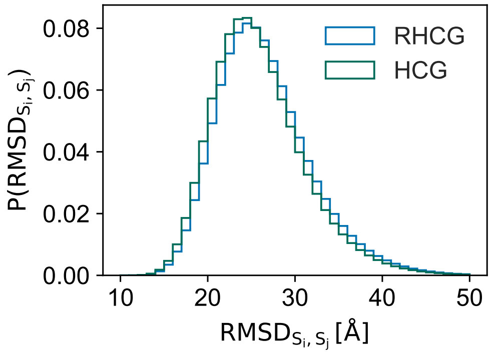


Figure S3: Conformational diversity of the tau K18 ensembles structures from HCG (green) and RHCG (blue). Shown is the distribution of the root-mean-squared distances (RMSD) of the $C_\alpha$ backbone atoms of two randomly chosen members of ensembles of size 5000.

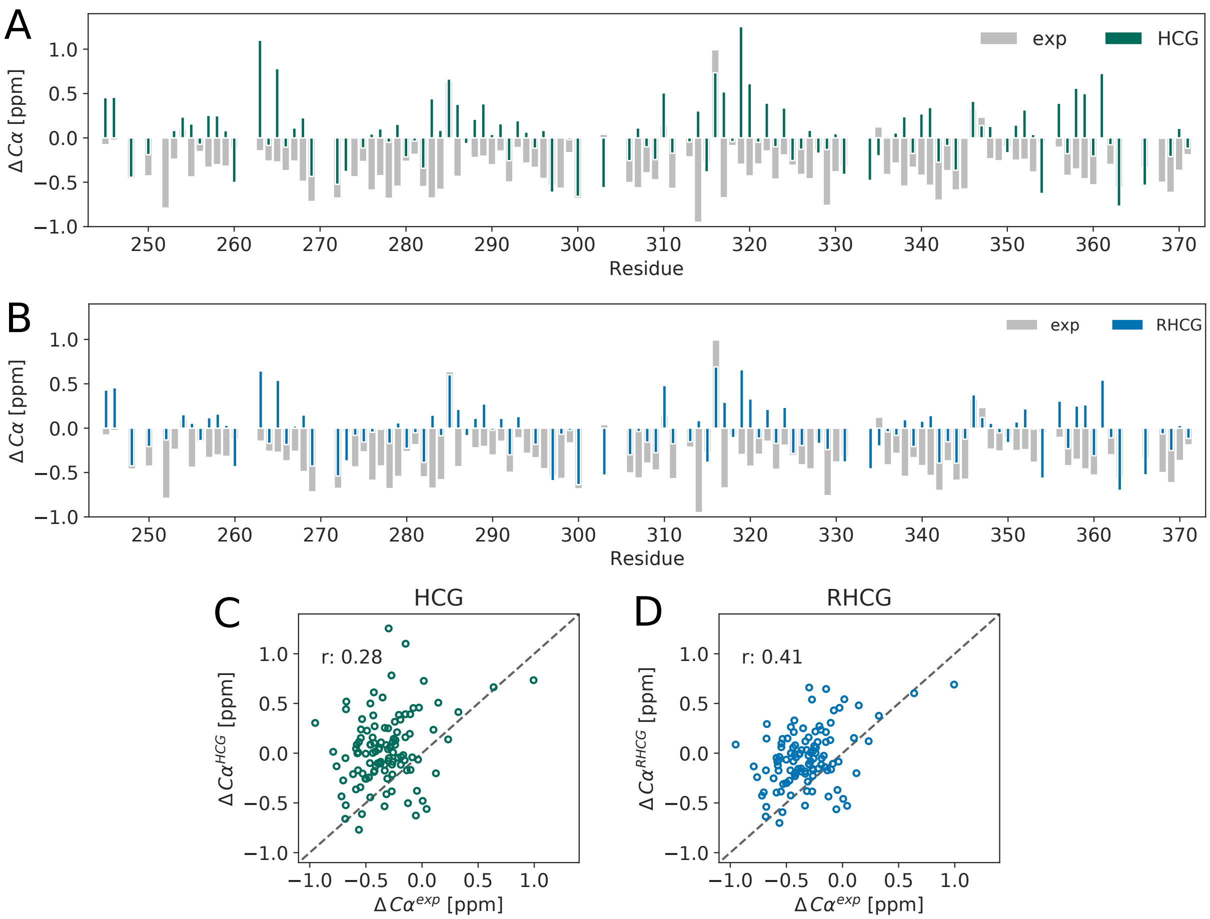


Figure S4: Comparison of calculated and measured NMR $C_\alpha$ chemical shifts. Experimental secondary shifts $\Delta C$ (grey bars) are compared as a function of residue number to the shifts calculated for the tau K18 ensembles from HCG (A, green bars) and RHCG (B, blue bars). (C,D) Scatter plot of experimental and calculated $\Delta C$ for HCG (C) and RHCG ensembles (D).

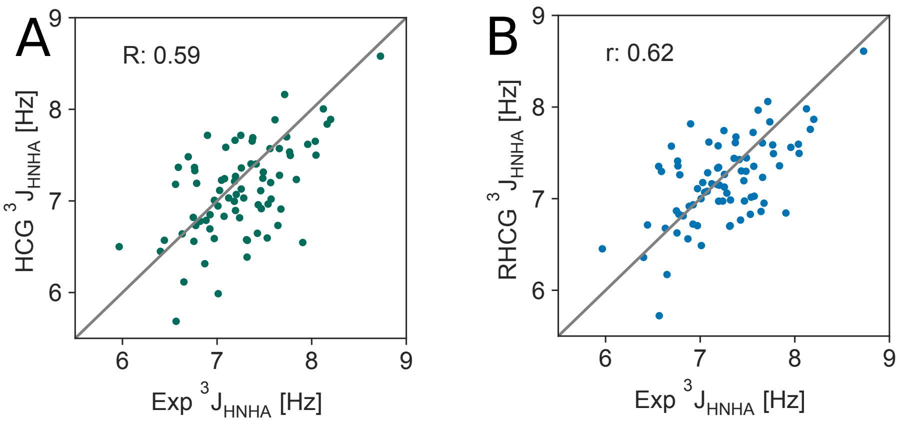


Figure S5: Scatter plot of $^3J_{\mathrm{HNH}\alpha}$ couplings computed from the structural ensembles created with (A) HCG and (B) RHCG against the couplings measured by NMR by Mukrasch et al.[S8]

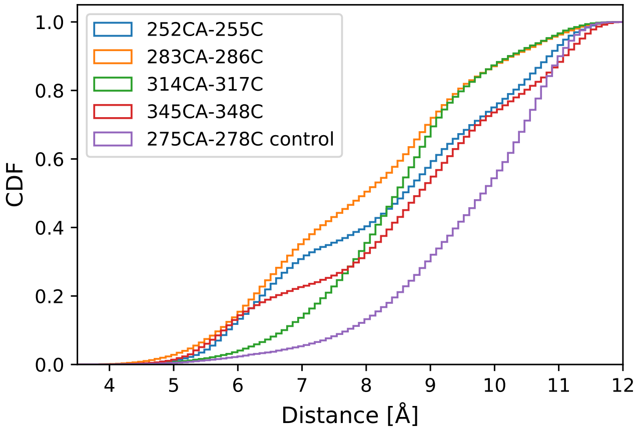


Figure S6: Cumulative distribution of $C_\alpha(n) - C'(n+3)$ distances for $^{252}$DLKN$^{255}$, $^{283}$DLSN$^{286}$, $^{314}$DLSK$^{317}$ and $^{345}$DFKD$^{348}$ in the RHCG ensemble. As control, the distance distribution for V275-I278 is shown (purple).

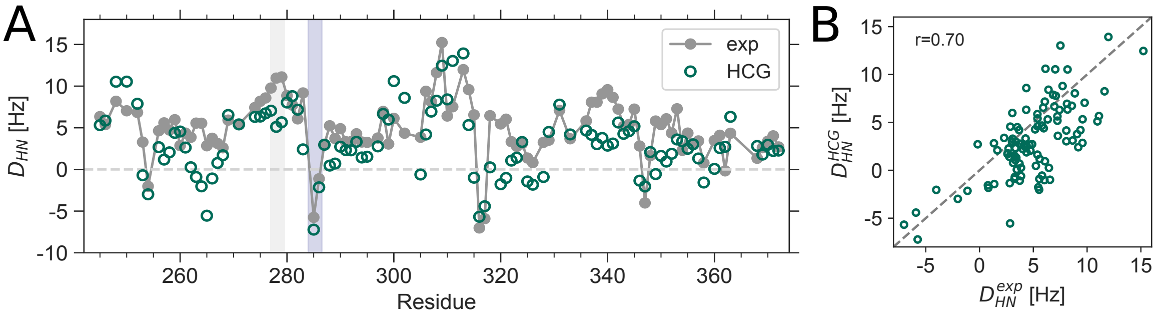


Figure S7: Comparison to NMR RDCs. (A) Experimental RDCs along the tau K18 sequence (grey) are compared to a tau K18 ensemble with 50 000 structures from HCG (dark green). (B) Scatter plot of RDCs calculated for the HCG ensemble and from experiment. The RDCs were calculated using PALES.[S22]

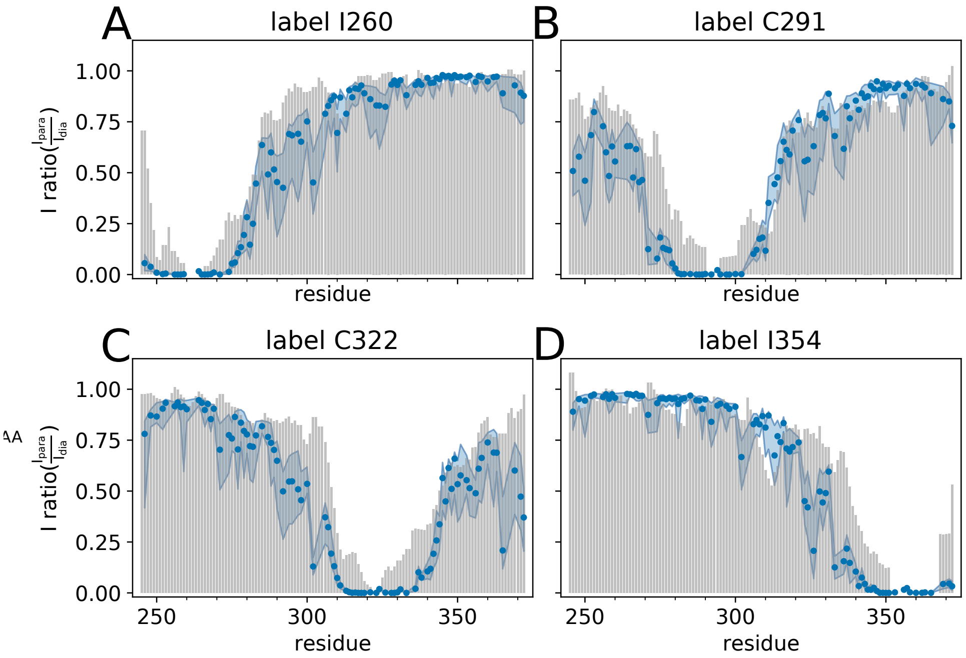


Figure S8: Paramagnetic relaxation enhancement analysis of tau K18. PRE profiles predicted for a large ensemble of tau K18 models from RHCG (blue dots) compared to the experimental profiles (grey bars). The blue dots indicate the median of the PREs predicted for 20 independent ensembles with 50 000 models each and the blue shaded area shows the interquartile range. PREs were predicted using PREpredict.[S9] The experimental PREs were measured at 278 K in dilute solution of tau K18. S-(1-oxyl-2,2,5,5-tetramethyl-2,5-dihydro-1H-pyrrol-3-yl)methyl methanesulfonothioate (MTSL) paramagnetic labels were attached to residues C260, I291, I322,and C354.

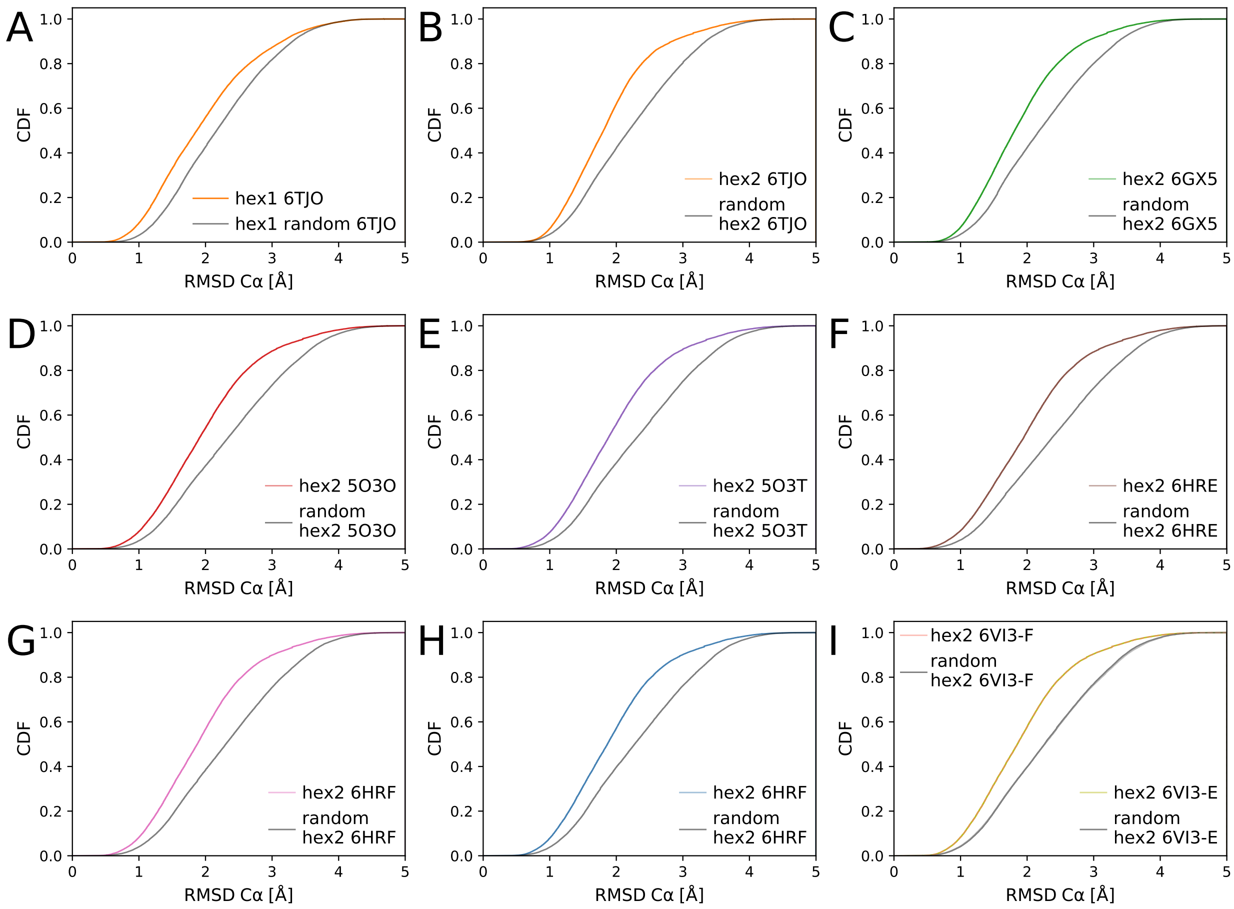


Figure S9: Comparison of hexapeptide structures from the RHCG ensemble to experimental structures of tau fibrils. Shown are cumulative distributions of the $C_\alpha$ RMSD of the first and second hexapeptide motif $^{275}$VQIINK$^{280}$ (A) and $^{306}$VQIVYK$^{311}$ (B-I), respectively. The PDB IDs of the different experimental reference structures are listed in the legends. For reference, gray lines show RMSD distributions for hexapeptide segments randomly chosen along the RHCG chains.

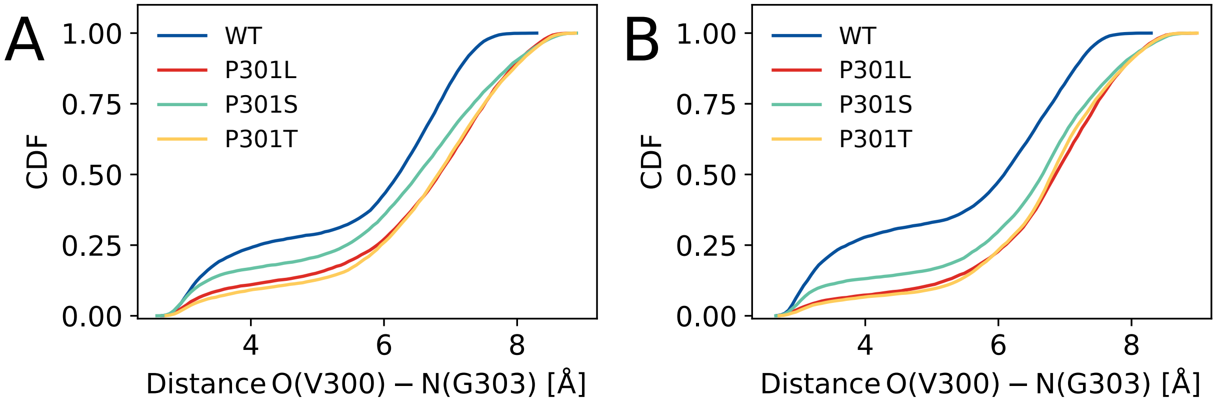


Figure S10: P301 mutants are locally more extended than wild-type tau K18. Cumulative distribution functions of O(V300)-N(G303) distances are shown for pentamer fragments with residue 301 at the (A) second and (B) central position for WT (blue) and P301L (red), P301S (turquoise), and P301T variants (yellow).

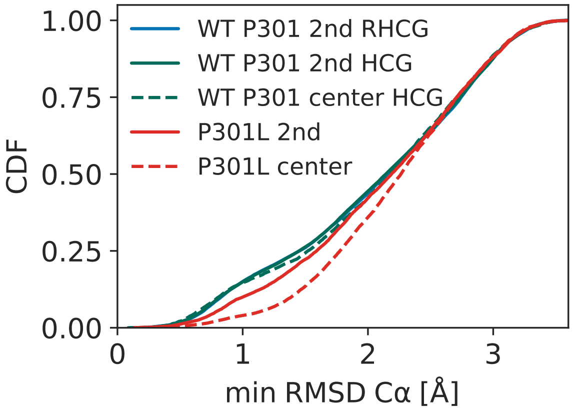


Figure S11: Comparison of the $^{300}$VPGGG$^{304}$ and $^{300}$VLGGG$^{304}$ motifs in the RHCG and HCG ensembles to NMR structures of microtubule-bound tau.[S13] Shown is the cumulative distribution function for the RMSD of the five $C_\alpha$ atoms with respect to the closest structure in the NMR ensemble (PDB: 2MZ7[S13]). Curves with solid and dashed lines correspond to ensembles created from fragments with the 301 position at the second and center position, respectively.
